# Genetic determinants of virulence and extensive drug resistance in *Pseudomonas aeruginosa* PPA14 isolated from eggplant rhizosphere

**DOI:** 10.1101/2023.06.03.543547

**Authors:** Sakthivel Ambreetha, Govindasamy Parshatd, Christian Castellanos, Giri Narasimhan, Dananjeyan Balachandar, Trevor Cickovski, Kalai Mathee

## Abstract

*Pseudomonas aeruginosa* is one of the Priority Level I critical pathogens that are least sensitive to antibiotics and can cause fatal hospital-acquired infections. This bacterium is predominantly present in the agricultural ecosystem. However, there are very limited studies on health threats associated with *P. aeruginosa* strains flourishing in edible plants. Previously, we isolated and characterized 18 *P. aeruginosa* strains from vegetable plants directly harvested from the farms. In the current work, it has been hypothesized that plant-associated *P. aeruginosa* harbors genetic determinants for virulence and resistance. To test this hypothesis, *in vitro* resistome profiles of the plant-associated *P. aeruginosa* strains were assessed based on the Kirby-Bauer disk diffusion method. Hierarchical clustering analysis was done to identify the plant-associated strains that are phenotypically similar to clinical isolates. An eggplant-associated strain, PPA14, that exhibited high virulence and extensive *in vitro* resistance against eight antibiotic classes was selected for complete genome analyses. The PPA14 genome was sequenced using the Solexa-Illumina and Oxford-Nanopore platforms, assembled, and annotated. The presence of virulence-related and antibiotic resistance (ABR) genes were predicted using the ABRicate tool and validated based on standard reference databases such as VFDB, NCBI AMRFinderPlus, MEGARes, CARD, and ResFinder. IslandViewer4 tool was used to predict the genes acquired through horizontal gene transfer. Additionally, comparative analyses of all the plant-associated and environmental *P. aeruginosa* genomes characterized so far were done using the Roary tool. The PPA14 genome size was 6.72 Mbp, encoding 6315 open reading frames. The genome harbored 49 ABR genes, including those coding for multiple families of efflux pumps that collectively confer resistance against 11 antibiotic classes. In addition, we detected 225 virulence-related genes, 83 genomic islands, and 235 unique genes in the PPA14 genome. Over 4% of the PPA14 genome is devoted to conferring virulence and extensive drug resistance. Our report highlights the health threat associated with an eggplant-associated *P. aeruginosa*.

## Introduction

Antibiotic-resistant (ABR) pathogens are emerging threats to the global health system. The World Health Organization has recorded 7,00,000 annual deaths due to ABR pathogens (1). *Pseudomonas aeruginosa* is one of the dreadful ESCAPE (*Enterococcus faecium, Staphylococcus aureus, Klebsiella pneumoniae, Acinetobacter baumannii, Pseudomonas aeruginosa, and Enterobacter species*) pathogens that cannot be eliminated by common antibiotics (2). In specific, carbapenem-resistant *P. aeruginosa* is a Priority Level-I Critical Pathogen that has been declared a serious threat (1, 3). *P. aeruginosa* has become a leading cause of life-threatening hospital-acquired infections due to its multi-drug resistant (MDR) phenotype. The MDR *P. aeruginosa*-associated infections can increase the fatality rate by 18 to 61% (4–6). Unfortunately, the MDR *P. aeruginosa* has been detected in diverse ecosystems. Improper disposal of hospital wastes potentially discharges several MDR *P. aeruginosa* into water bodies (7). Numerous MDR *P. aeruginosa* strains have been identified in wastewater effluents, polluted water bodies, hydrocarbon-contaminated soil, and compost (8–13). Contaminated water channels and farm animals have been identified as the huge reservoirs of the MDR *P. aeruginosa* strains (14, 15). These strains could be easily transmitted to agricultural fields and plants.

In our previous studies, we isolated and characterized 18 *P. aeruginosa* strains from the rhizospheric and endophytic niches of edible plants (cucumber, tomato, eggplant, and chili) harvested directly from the farms (16, 17). DNA fingerprinting and 16S rDNA-based phylogenetic analyses showed that these strains are evolutionarily related to the clinical *P. aeruginosa* isolates (16). Nearly 50-80% of these plant-associated *P. aeruginosa* strains exhibited multiple virulence traits including lytic activities (hemolysis, proteolysis, and lipolysis), biofilm formation, swarming motility, and production of rhamnolipids, pyocyanin, and siderophores (16, 17). Also, 10-40% of death occurred in the *C. elegans* model when the *P. aeruginosa* strains from the agricultural system were given as the food source (17). These reports indicate the presence of virulent and pathogenic *P. aeruginosa* strains in edible plants.

Furthermore, *P. aeruginosa* contamination has been detected in fresh vegetables at more than 50% of the supermarkets (18). Contaminated agricultural produces could be a major source of human transmission of *P. aeruginosa* (19, 20). Since *P. aeruginosa* is transmittable across the plant-animal-human interface (19–26), it is important to analyze the genetic determinants of virulence and pathogenicity in the non-clinical strains. However, the genotypic characteristics of plant-associated *P. aeruginosa* strains have not been studied as extensively as that of clinical isolates. Only six out of 214 *P. aeruginosa* complete genomes available in the *Pseudomonas* database belong to the plant-associated strains (https://pseudomonas.com/). There is a clear gap in the genomic characterization of the agricultural *P. aeruginosa* strains.

The omnipresence of *P. aeruginosa* is often supported by a large and complex genome (5.5 to 7.4 Mbp; (27)). *P. aeruginosa* flourishes in diverse habitats by frequently acquiring new genetic elements or genomic islands through horizontal gene transfer (HGT) (28–31). Nearly 3000 plasmids potentially acquired through HGT have been identified in various *P. aeruginosa* strains (31). As a result, this bacterium has evolved with a highly versatile genome. Only 6.6% of its genome comprises core essential genes while the rest could be variable among the strains (32). Such variable regions in the *P. aeruginosa* genome contribute to the strain-level metabolic and functional diversity and are termed the regions of genomic plasticity (28, 33, 34). Importantly, 5 to 12% of the variable regions harbor virulence-related and drug-resistance genes (31). Previous reports suggest that the pathogenicity of *P. aeruginosa* is often related to genomic flexibility (35, 36). Henceforth, complete genome analyses of plant-associated *P. aeruginosa* strains might help to identify if they harbor genetic elements that confer virulence, pathogenicity, and ABR.

In the current study, we looked into the *in vitro* ABR profile of 18 plant-associated *P. aeruginosa* strains. An eggplant rhizospheric strain, PPA14 that exhibited extensive resistance was selected for complete genome analyses. It was hypothesized that plant-associated *P. aeruginosa* (PPA) acquires virulence-related and drug-resistance genes from the environment. To test this hypothesis, we examined the genomic islands, virulence-related, and ABR genes in the PPA14 genome. In addition, the PPA14 genome was compared with other plant-associated and environmental *P. aeruginosa* strains to determine the unique and shared genes among the non-clinical isolates.

## Methods

### Bacterial strains used in this study

Eighteen plant-associated *P. aeruginosa* strains (PPA01-18) used in this study (Table 1) were previously isolated and characterized from vegetable plants such as cucumber, tomato, eggplant, and chili (16, 17). Clinical strains of *P. aeruginosa*, PAO1, ATCC10145, and ATCC9027 were used as controls for phenotypic assays (37–39). The *P. aeruginosa* PA14 genome was used as the reference for genome assembly (40). The complete genomes of six plant-associated strains and 16 environmental strains of *P. aeruginosa* that were previously sequenced and deposited in the *Pseudomonas* database (https://pseudomonas.com/) were used for comparative genomic analyses along with the standard reference genomes, PAO1 and PA14 (40, 41).

**Table 1.**
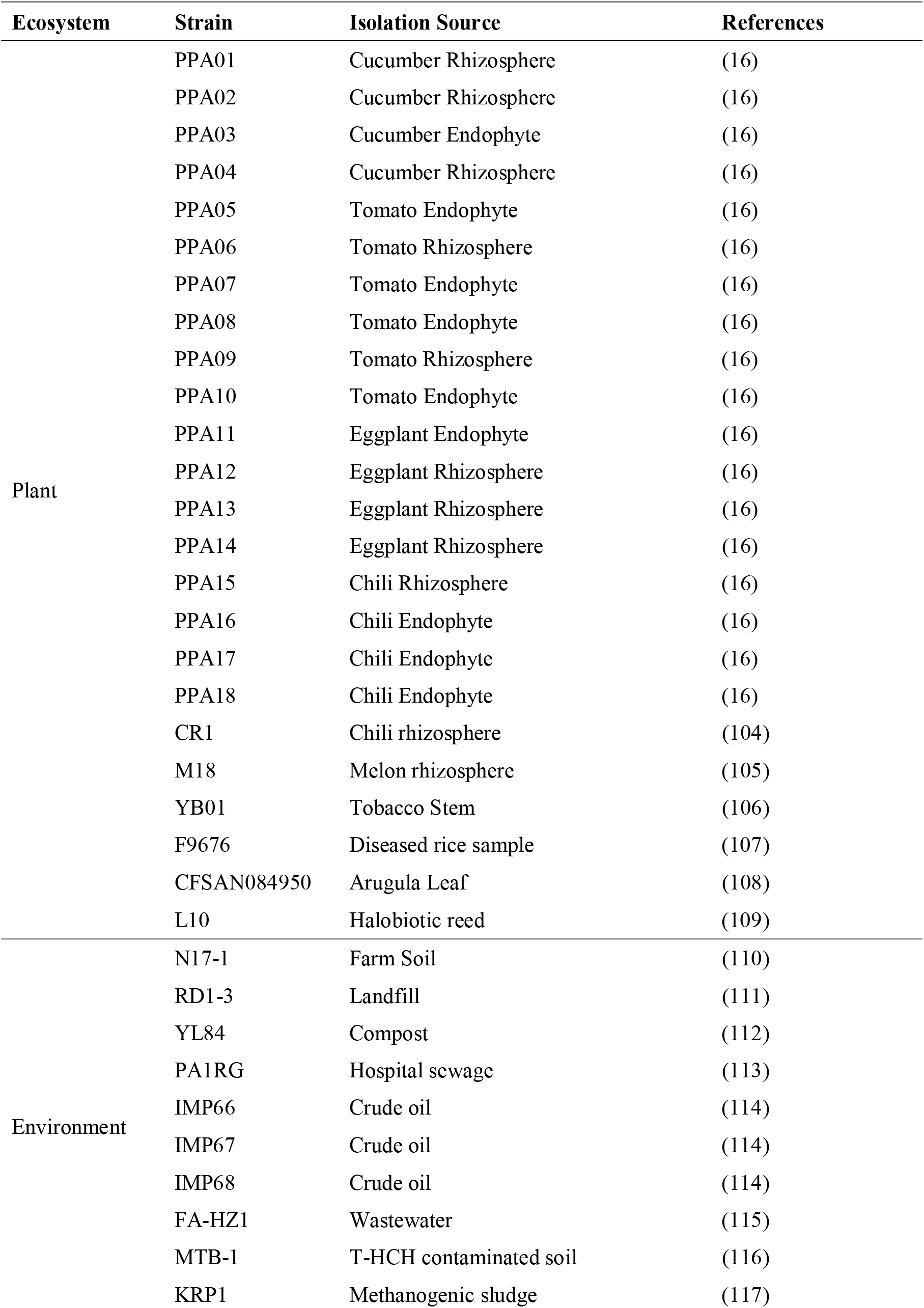
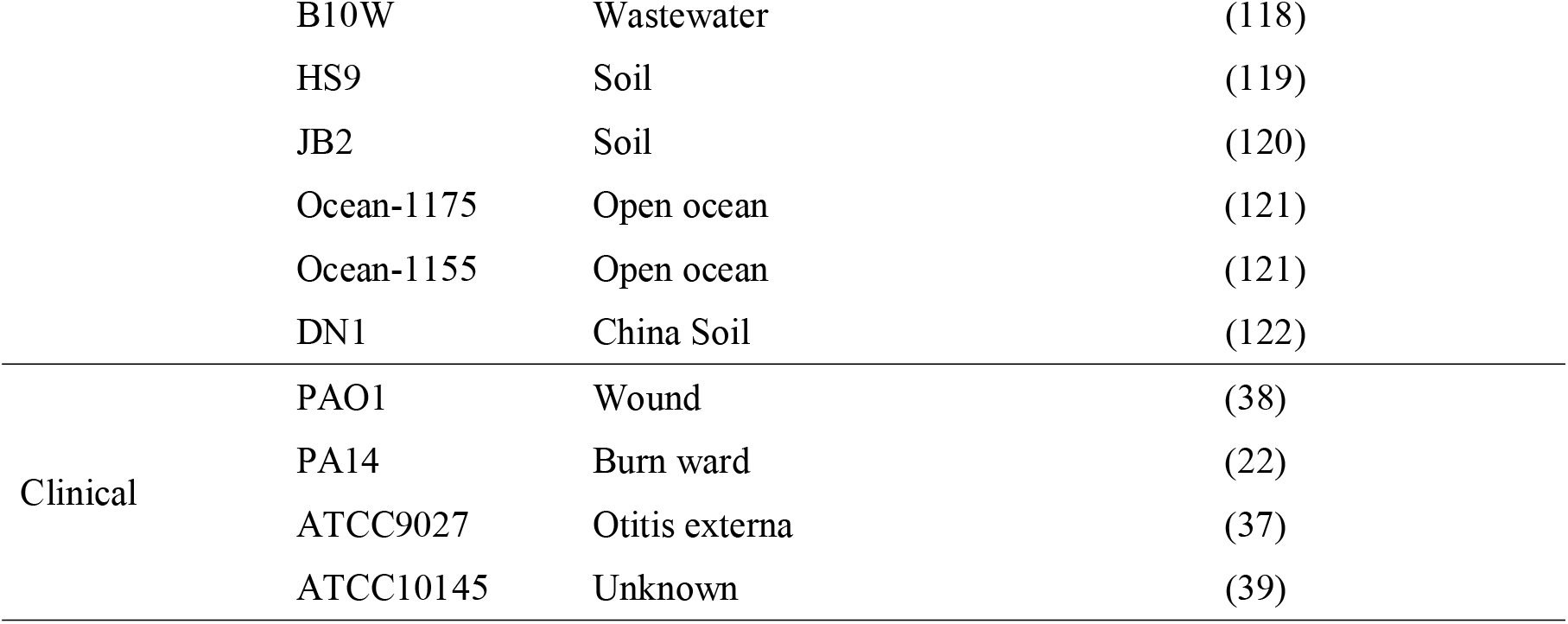
*Pseudomonas aeruginosa* strains used in this study.

### Antibiotic resistance assay

The ABR profiles of the *P. aeruginosa* strains were tested using the Kirby-Bauer disk diffusion method (42, 43). The three clinical strains of *P. aeruginosa*, ATCC10145, ATCC9027, and PAO1 were used as controls. Antibiotic susceptibility test was done using octa- and dodeca discs (Himedia) with 18 different antibiotics from eight classes namely aminoglycosides (gentamycin, streptomycin, and kanamycin), fluoroquinolone (ciprofloxacin), cephalosporins of first (cefadroxil), second (cefuroxime, and cefaclor), and third (cefotaxime, and cefoperazone) generations, penicillins (ampicillin, augmentin, and penicillin-G), macrolides (erythromycin, azithromycin, and clarithromycin), nitrofuran (nitrofurantoin), polymyxin (colistin), sulphonamide (co-trimoxazole), and tetracycline. Briefly, the strains were grown overnight at 37°C in Luria Bertani (LB) broth. After 24 h, 10% of the OD_600_∼1.0 adjusted overnight cultures were added into 15 ml of LB medium at a lukewarm temperature and poured into sterile Petri plates. After solidification, the antibiotic discs were placed over the inoculated media, and the plates were incubated overnight at 37°C. The occurrence of the inhibition zone around a particular disc was indicative of susceptibility toward the corresponding antibiotic. The percentage of antibiotic resistance exhibited by each of the tested strains and the percentage of strains resistant to each of the tested antibiotics were computed.

### Hierarchical clustering of *P. aeruginosa* strains

The phenotypic and metabolic traits of the 18 PPA strains and the three clinical strains, ATCC10145, ATCC9027, and PAO1 were characterized previously (16, 17). In the current study, all these *P. aeruginosa* strains were grouped based on their phenotypic relatedness using the NCSS 2020 Statistical Software (https://www.ncss.com/software/ncss/). A hierarchical-clustered heat map with a double dendrogram was generated based on the Manhattan distance metric and the similarity between the clusters was defined using Ward’s minimum variance (44). The heat map and the dendrogram were used to identify the most virulent plant-associated strains that are closely related to the clinical strains.

### Extraction and quantification of *P. aeruginosa* PPA14 genomic DNA

The genomic DNA of *P. aeruginosa* PPA14 was extracted using the hexadecyl-trimethyl ammonium bromide (CTAB) method (45) and quantified using the NanoDrop (Thermo Scientific, Nanodrop 2000c) spectrophotometer. The absorbance relation of the DNA at 260/280 was indicative of its purity.

### Whole-genome sequencing

The complete genomic DNA of the *P. aeruginosa* PPA14 strain was sequenced using the Solexa-Illumina, and Oxford-Nanopore technologies platforms, for short and long reads, respectively (46–48) by the Genotypic Technology, Bangalore, India (www.genotypic.co.in).

### Genome assembly and scaffolding

The *P. aeruginosa* PPA14 genome reads were assembled using eleven different software tools (Fig. 1; (49)). The short reads generated by the Solexa-Illumina sequencing were assembled using SPAdes v3.13.0 (50), IDBA v1.1.3 (51), and Megahit v1.1.3 (52), SKESA v2.3.039 (53), and Unicycler v0.4.711 (54). The long reads generated by the Oxford-Nanopore technologies were assembled using Canu 1.8 (55), Flye 2.3.7 (56), and Unicycler v0.4.7 (54). Unicycler v0.4.7 (54), SPAdes v3.13.0 (57), and IDBA-hyb v1.1.1 (51) were used to assemble the hybrid reads formed by the combination of short and long reads. The contig level assemblies generated by each of these tools were further assembled into scaffolds using MeDuSa v1.6 (58) with *P. aeruginosa* PA14 as the reference genome.

**Fig. 1.**
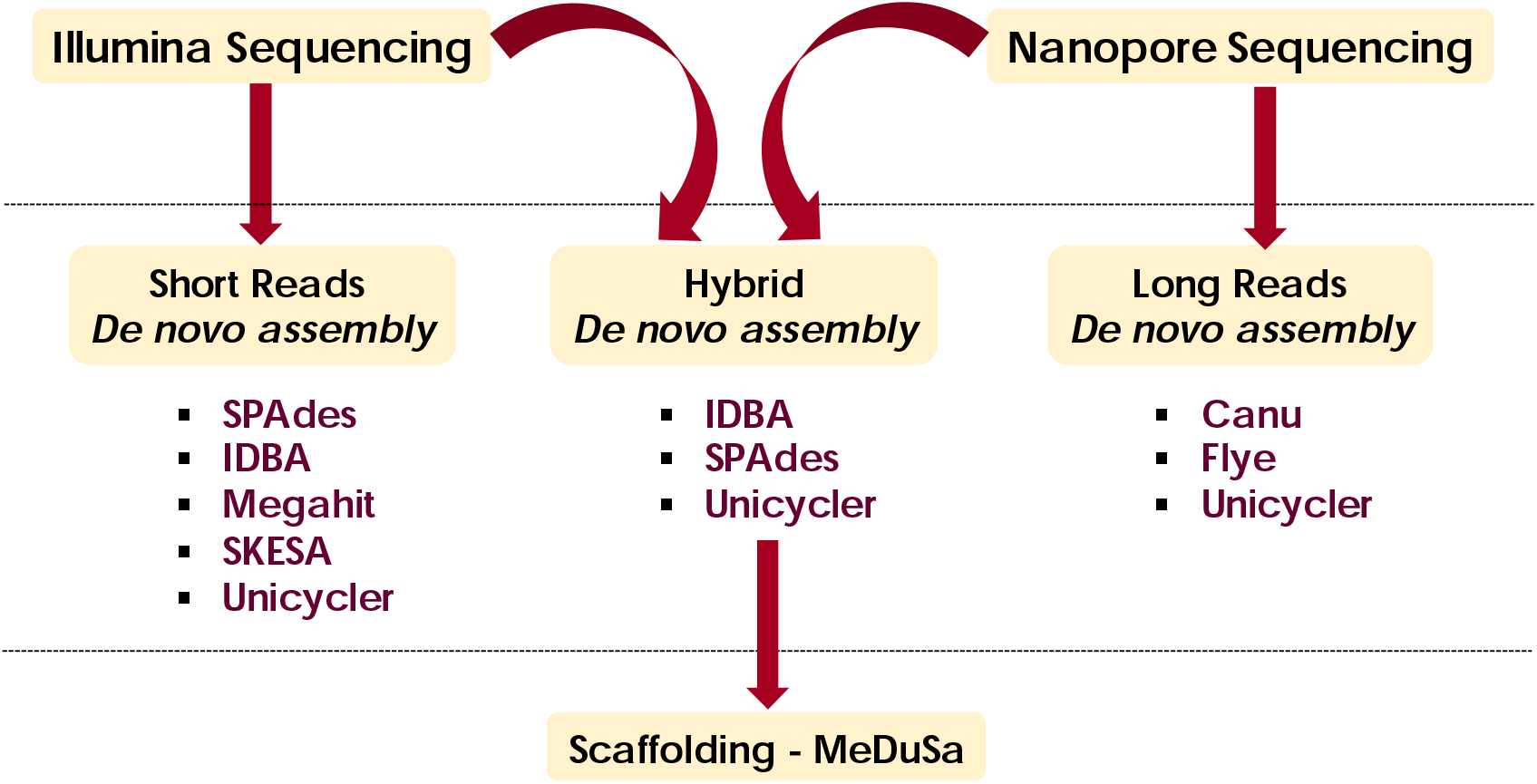
*Pseudomonas aeruginosa* PPA14 genome sequencing and assembly. The flow chart represents the different sequencing platforms and assembly tools used for the sequencing and assembly of the PPA14 genome (Method adapted from Molina-Mora et al. 2020). SPAdes v3.13.0, IDBA v1.1.3. and Megahit v1.1.3, SKESA v2.3.039, and Unicycler v0.4.711, Canu 1.8, and Flye 2.3.7 were the software versions used.

### Comparison and evaluation of genomic assemblies

All 11 genome assemblies were compared using four different tools, QUAST 5.0.1 (59), BUSCO (60), Circlator (61), and Qualimap v2.2.2 (62); Fig. 2). The assemblies were evaluated based on their genome and annotation features such as number of scaffolds, genome size, length of the shortest contig that covers 50% of the genome size (N50), orthologous gene score, number of repeat regions, and number of rRNAs and tRNAs (49). The best-assembled genome was selected based on these metrics and used for further downstream analyses.

**Fig. 2.**
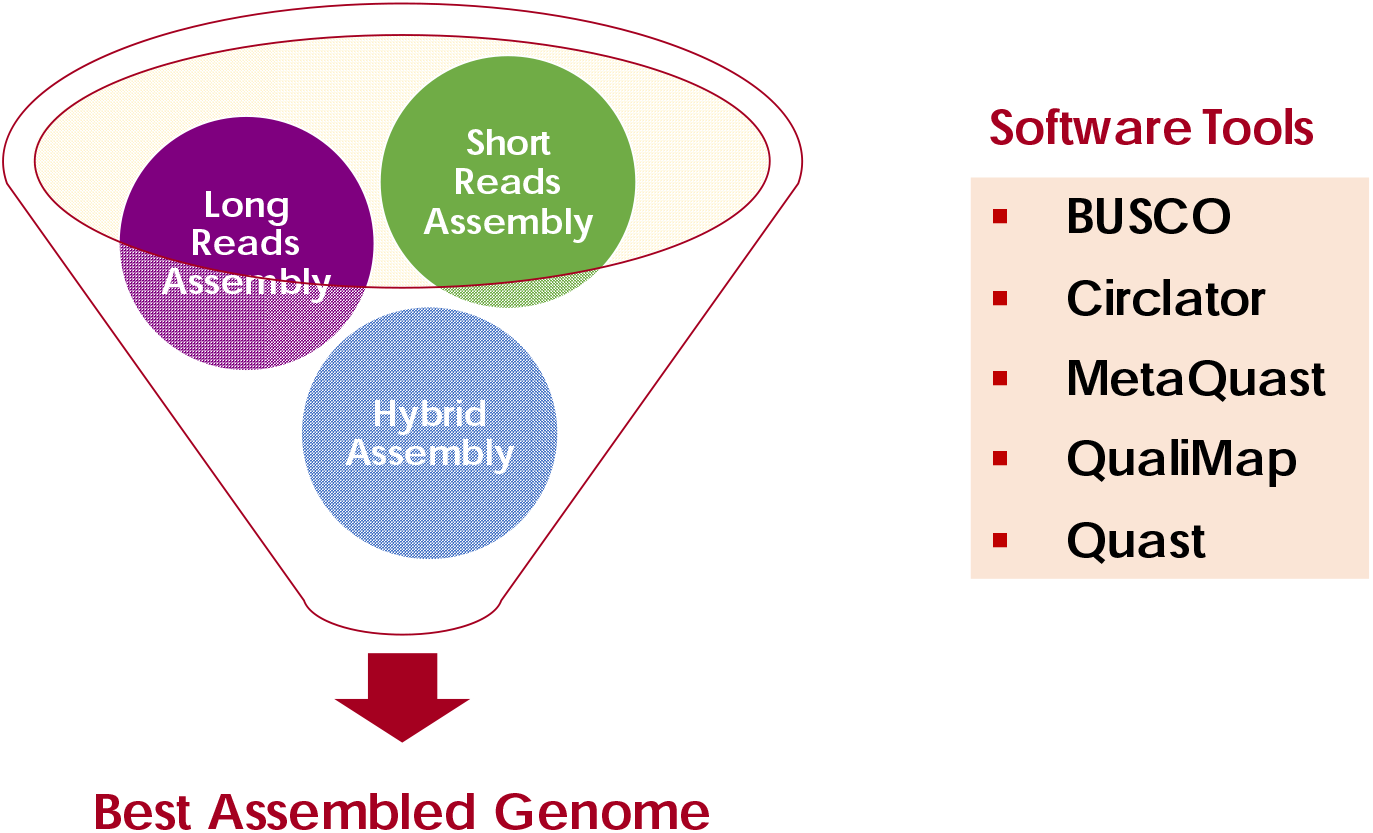
Comparison and evaluation of *P. aeruginosa* PPA14 genome assemblies. The figure represents the list of tools used to select the best-assembled draft genome by screening all the short, long, and hybrid assemblies of the PPA14 genome (Method adapted from Molina-Mora et al. 2020).

### Gene annotation

The assembled genome of PPA14 was annotated using the rapid prokaryotic genome annotation (Prokka) and Rapid annotation using subsystem technology (RAST) computational tools to predict its functional motifs (63–65). Furthermore, the predicted gene functions were manually curated using the standard *P. aeruginosa* PAO1, PA14, PAK, and PA7 genomes as the references.

### Identification of ABR and virulence genes

The ABRicate tool (https://github.com/tseemann/abricate) was used to identify the genetic determinants of *P. aeruginosa* PPA14 drug resistance and virulence. The AMR genes were detected based on the AMRFinderPlus, National Center for Biotechnology Information (NCBI), comprehensive antibiotic resistance database (CARD; (66)), and ResFinder, Center for Genomic Epidemiology (CGE; (67)). The virulence genes were identified based on the virulence factor database (VFDB; http://www.mgc.ac.cn/VFs/; (68)).

### Identification of genomic islands

The Island Viewer 4 tool was used to identify the genomic islands (GI) harbored by the *P. aeruginosa* PPA14 genome (69). The Island Viewer 4 tool predicted the GIs in the PPA14 genome using the ensemble method by integrating three different tools, IslandPath-DIMOB, SIGI-HMM, and IslandPick (69). IslandPath-DIMOB detects the GIs based on the presence of at least one mobile genetic element coupled with dinucleotide bias in more than eight consecutive genes (70). SIGI-HMM predicts the GIs based on the hidden Markov model depending on biased codon usage (71, 72). IslandPick uses a comparative genomics approach on monophyletic strains to identify the GIs (73).

### Genome map

The genome map of *P. aeruginosa* PPA14 was created and visualized using the circular genome viewer (CGView) server ((74); http://cgview.ca/). The open reading frames (ORFs), functional motifs, GC content, GC bias, predicted GIs and ABR genes were integrated and visualized in the PPA14 genome map.

### Comparative genomic analyses of agricultural, and environmental *P. aeruginosa* strains

Complete genomes of *P. aeruginosa* strains that were previously isolated from plants (CR1, YB01, F9676, M18, CFSAN084950, and PPA14) and environments (N17-1, RD1-3, YL84, IMP66, IMP67, IMP68, PA1RG, MTB-1, KRP1, B10W, JB2, Ocean-1175, Ocean-1155, HS9, FA-HZ1, and DN1) were downloaded from the *Pseudomonas* database for the comparative genomic analyses (75). Two reference *P. aeruginosa* genomes, PAO1 and PA14, were included as controls (Table 1). Computational tools such as Prokka (65), Roary, the pan-genome pipeline (76), and FastTree (77) were used for comparative genomic analyses of the 25 *P. aeruginosa* strains. The pan-genome profile of these strains produced based on the ‘gene presence-absence matrix’ produced by the Roary tool was visualized using Phandango (https://jameshadfield.github.io/phandango/<u>#/</u>; (78)). The unique genes harbored by the PPA14 genome were identified from the pan-genome profile and their functions were predicted by BLAST analyses against the reference genomes in the NCBI database. The maximum composite likelihood method-based phylogenetic analysis was performed using the complete genome of the 25 *P. aeruginosa* strains analyzed in this study. The phylogenetic tree was constructed using the randomized axelerated maximum likelihood (RAxML) computation tool (79).

### Statistical analyses

All the phenotypic experiments were performed in triplicates. All data were subjected to a one-way analysis of variance (ANOVA) with a *p*-value of 0.05, and Duncan’s multiple range test (DMRT) was performed between individual means to reveal any significant difference (XLSTAT, version 2010.5.05 add-in with Windows Excel). The hierarchical-clustered heat map with a double dendrogram was created using NCSS 2020 statistical software (NCSS, Kaysville, USA) to cluster the *P. aeruginosa* strains based on their phenotypic characteristics. Data analysis and scientific graphing were done in OriginPro version 8.5 (OriginLab®, USA).

## Results

### Phenotypic drug-resistance profile of *P. aeruginosa* strains

*In vitro* ABR profile of the plant-associated and clinical *P. aeruginosa* strains was assessed using 19 antibiotics belonging to eight different classes including aminoglycosides, beta-lactams, tetracyclines, fluoroquinolones, macrolides, nitrofurans, polymyxin, and sulfonamides (list of antibiotics is given in the “Materials and Methods”).

The efficacy of different antibiotics in curtailing *P. aeruginosa* growth was estimated based on the percentage of strains resistant to a particular antibiotic (Fig. 3A). All of the plant-associated *P. aeruginosa* strains were inhibited by a fluoroquinolone (10 µg/ml ciprofloxacin). Cefotaxime (30 µg/ml), colistin (10 µg/ml), and gentamycin (10 µg/ml) belonging to third-generation cephalosporin, polymyxin, and aminoglycoside classes, respectively were effective against 84-95% of the PPA strains. A macrolide (15 µg/ml erythromycin), and a second-generation cephalosporin (30 µg/ml cefaclor), prevented the growth of 77 to 88% of the PPA strains. Only 10% of the tested strains were sensitive to first-generation cephalosporins (30 µg/ml cefadroxil), nitrofuran (300 µg/ml nitrofurantoin), and sulphonamide (25 µg/ml co-trimaxazole). The rest of the antibiotics exhibited an average resistance and inhibited about 30 to 60% of the PPA strains tested in this study (Fig. 3A).

**Fig 3.**
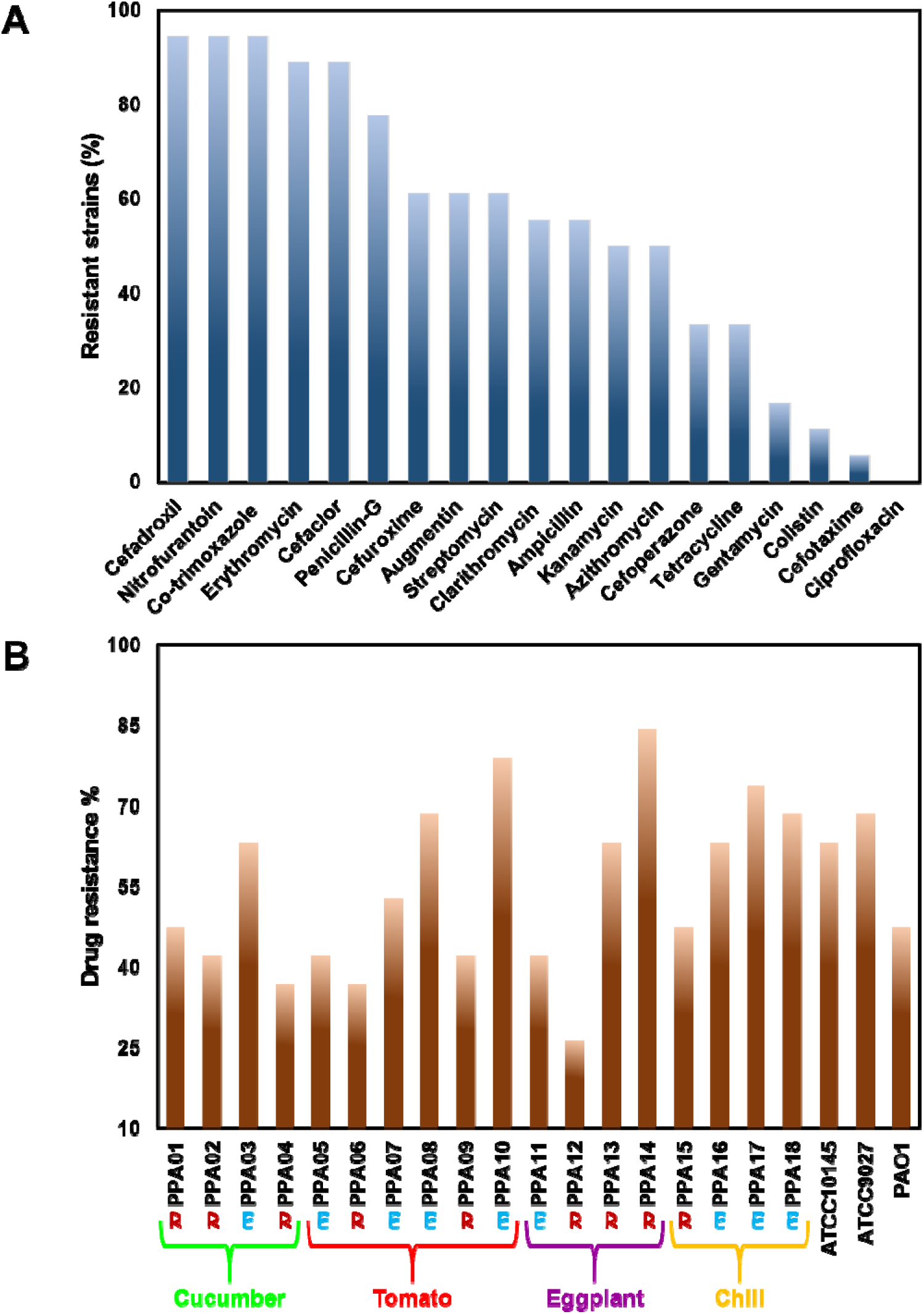
Antibiotic resistance profile of *P. aeruginosa* strains. (A) The efficacy of different antibiotics in inhibiting plant-associated *P. aeruginosa* strains. (B) The percentage of drug resistance exhibited by each of the tested plant-associated and clinical strains of *P. aeruginosa.* The strains are color-coded based on their source of isolation. Green – cucumber; red – tomato; purple – eggplant; yellow – chili; black – clinical isolates.

*In vitro* resistance profile of each *P. aeruginosa* strain was recorded individually (Fig. 3B). The three clinical isolates (ATCC10145, ATCC9027, and PAO1) were resistant against 47 to 68% of the antibiotics. None of the plant-associated *P. aeruginosa* strains were susceptible to all the 19 antibiotics tested in this work. Importantly, 12 out of 18 PPA strains tested in this study had higher drug resistance than the *P. aeruginosa* PAO1. The overall resistance percentage of the plant-associated strains varied between 26 to 84%. The strains, PPA12, and PPA14 isolated from the eggplant rhizosphere exhibited the lowest and highest level of drug resistance, respectively. Seven out of the nine endophytic (PPA03/cucumber; PPA07, PPA08/tomato; PPA16, PPA17, PPA18/chili) and two out of the nine rhizospheric (PPA13, PPA14/eggplant) strains resisted more than 50% of the tested antibiotics. The rest of the strains (PPA01, PPA02, PPA04/cucumber; PPA05, PPA06, PPA09, PPA10/tomato; PPA11, PPA12/eggplant; PPA15/chili) exhibited relatively low antibiotic resistance.

### Phenotypic relatedness of agricultural and clinical *P. aeruginosa* strains

The plant-beneficial traits, virulence factors, and pathogenicity levels of plant-associated (PPA01-PPA18) and clinical (ATCC10145, ATCC9027, and PAO1) *P. aeruginosa* strains used in this study were previously determined (16, 17). This includes production of pyocyanin, rhamnolipid, siderophores, ammonia, and indole-3 acetic acid, formation of biofilm, swarming motility, solubilization of complex soil minerals, hemolytic, lipolytic, and proteolytic activities, biocontrol of phytopathogenic bacteria (*Xanthomonas oryzae*) and fungi (*Pythium aphanidermatum*, *Rhizactonia solani*, and *Fusarium oxysporum*), and pathogenicity against the *Caenorhabditis elegans* model (16, 17).

In the current work, a hierarchically-clustered heat map with a double dendrogram was created based on these phenotypic traits and drug-resistance profiles to identify the plant-associated *P. aeruginosa* strains that are closely related to the clinical isolates (Fig. 4). The cells in the heat map were color-coded from blue (low) to red (high) based on the relative abundance of the traits across the tested strains. All the tested strains clustered into three distinct groups (top dendrogram). Cluster A was occupied by seven strains (PPA01/cucumber; PPA05, PPA06, PPA07, PPA09/tomato; PPA11, PPA12/eggplant). Two cucumber-associated strains (PPA02 and PPA04) were found in cluster B. Cluster C had three clinical isolates (ATCC10145, AT9027, and PAO1) and nine plant-associated strains (PPA03/cucumber; PPA08, PPA10/tomato; PPA13, PPA14/eggplant; PPA15, PPA16, PPA17, PPA18/chili). These nine strains exhibited shared virulence, pathogenicity, and drug resistance with the tested clinical isolates. Among these plant-associated strains, PPA14 isolated from an eggplant rhizosphere caused 35% mortality in the *C. elegans* model (17) and exhibited extensive resistance against eight antibiotic classes (aminoglycosides, fluoroquinolone, three-generation cephalosporins, penicillins, macrolides, nitrofuran, sulphonamide, and tetracycline) used in the current work (Fig. 3). Since the *P. aeruginosa* PPA14 expressed relatively higher pathogenicity and drug-resistance among the 18 tested plant-associated strains and also got clustered with the clinical *P. aeruginosa* isolates (Fig. 4), this strain was selected for the complete genome analyses.

**Fig. 4.**
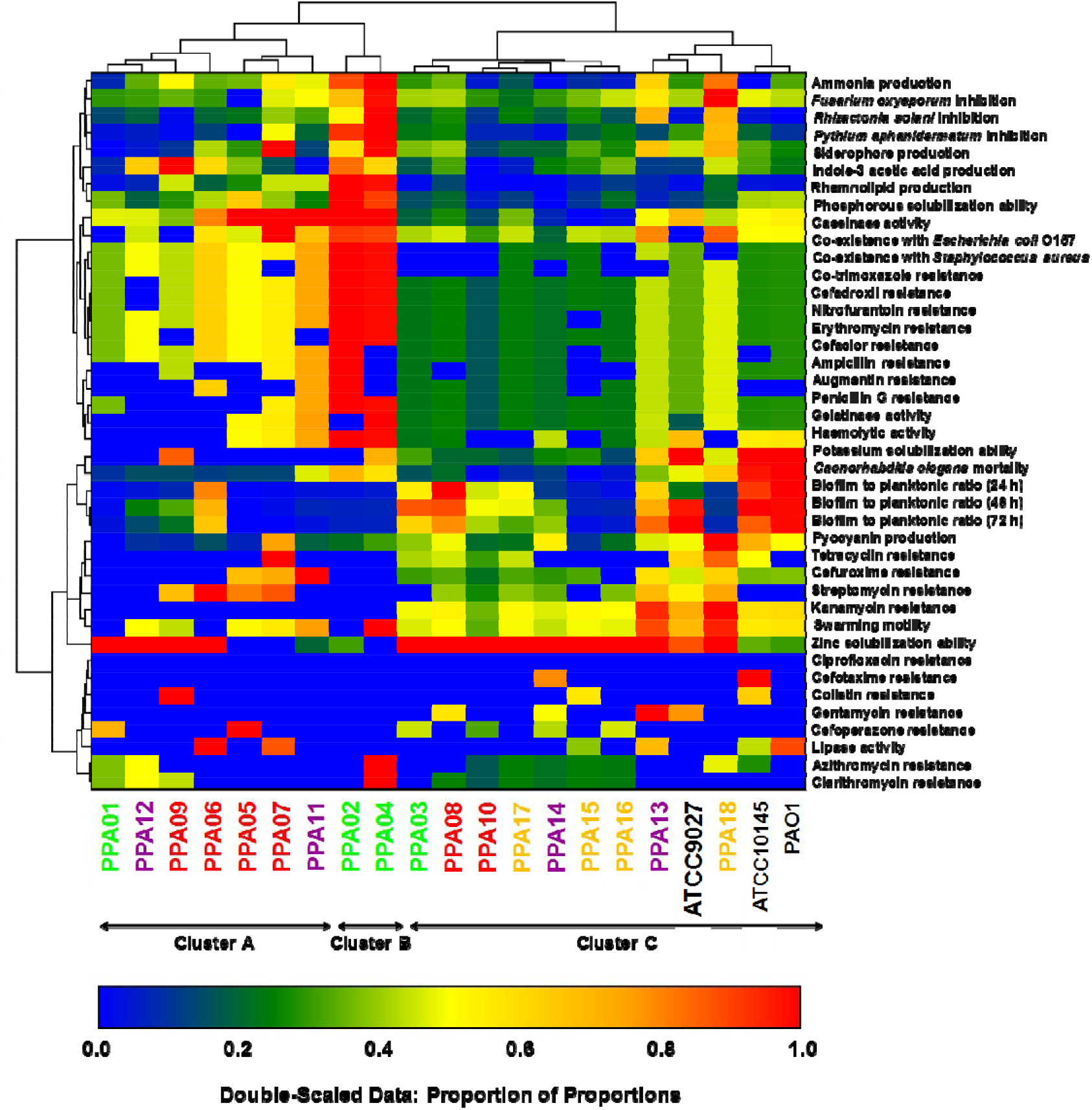
Hierarchical clustering analysis of *P. aeruginosa* strains. Double dendrograms (top and left) depict the relatedness between strains and between traits, respectively. Hierarchical clustering was done based on Ward minimum variance with Manhattan distance as the distance metric. The proportion method was used for variable scaling. Cell colors correspond to the relative abundance of the traits after mean-centering and unit-scaling, ranging from blue (low value) to red (high value). The strain names are color-coded based on their isolation source, green – cucumber, red – tomato, purple – eggplant, yellow – chili, black – clinical origin.

### *P. aeruginosa* PPA14 genome sequencing and assembly

The whole genome of *P. aeruginosa* PPA14 was sequenced using two platforms, Solexa-Illumina and Oxford-Nanopore technologies (46–48). The Illumina and Nanopore sequencing provided short and long raw reads, respectively. These reads were assembled individually and also combined to generate hybrid assemblies (refer to ‘Materials and Methods’ for the list of assembly tools). MeDuSa v1.6 (58) was used for scaffolding the assemblies based on *P. aeruginosa* PA14 as the reference genome. In total, 11 draft genome assemblies of PPA14 were generated which included five short-read, three long-read, and three hybrid-read assemblies. The 11 draft genome assemblies of the PPA14 were compared using QUAST 5.0.1 (59), BUSCO (60), Circlator (61), and Qualimap v2.2.2 (62). Each of these assemblies was evaluated based on their contiguity and completeness (49). The number of scaffolds generated by each assembly tool was considered the primary metric for contiguity. The short-assemblies had a higher number of scaffolds compared to long- and hybrid-assemblies (Fig. 5a). Only Hybrid-SPAdes, Hybrid-IDBA, Long-Canu, and Long-Unicycler could produce a single circular scaffold that reflected their contiguity. The short-assemblies also had low performance based on their i) draft genome length and ii) length of the shortest contig that covers 50% of the genome size (N50; Fig. 5b). The Long-Canu and Short-IDBA assemblies had the largest (6.74 Mbp) and shortest (6.59 Mbp) draft genome, respectively. The contiguity of an assembly is determined based on its N50 (49). In this study, the draft genome length was equal to the N50 only in the assemblies produced by the Hybrid-SPAdes, Hybrid-IDBA, Long-Canu, and Long-Unicycler. An orthologous gene score was generated for each of the assemblies based on the presence of conserved genes. The assemblies with more than 99.5% orthologous gene scores were considered to be the best (60). Three short-assemblies (Megahit, SKESA, and Unicycler) and two hybrid assemblies (IDBA, and Unicycler) had a 99.9% orthologous gene score (Fig. 6a). All three long assemblies had poor scores based on their orthologous genes (75 to 95%). The Short-Megahit (203) and Short-IDBA (39) had the highest and lowest number of repeat regions, respectively. Seven out of eleven assemblies (Short-SPAdes and SKESA; Long-Canu, Flye, and Unicycler; Hybrid-SPAdes and Unicycler) had nearly 100 repeat regions. The *P. aeruginosa* genomes harbor a minimum of 12 ribosomal RNAs (rRNAs) and 64 transfer RNAs (tRNAs). On comparing all the PPA14 genome assemblies generated in this study, the tRNA numbers ranged from 36 to 64 while the rRNA numbers ranged from 2 to 12. Only three out of eleven assembly tools, Long-Flye, Hybrid-SPAdes, and Hybrid-Unicycler generated genomes with the optimal number of RNAs (Fig. 6b). Particularly, the rRNA count was too low (>8) in the rest of the assemblies.

**Fig. 5.**
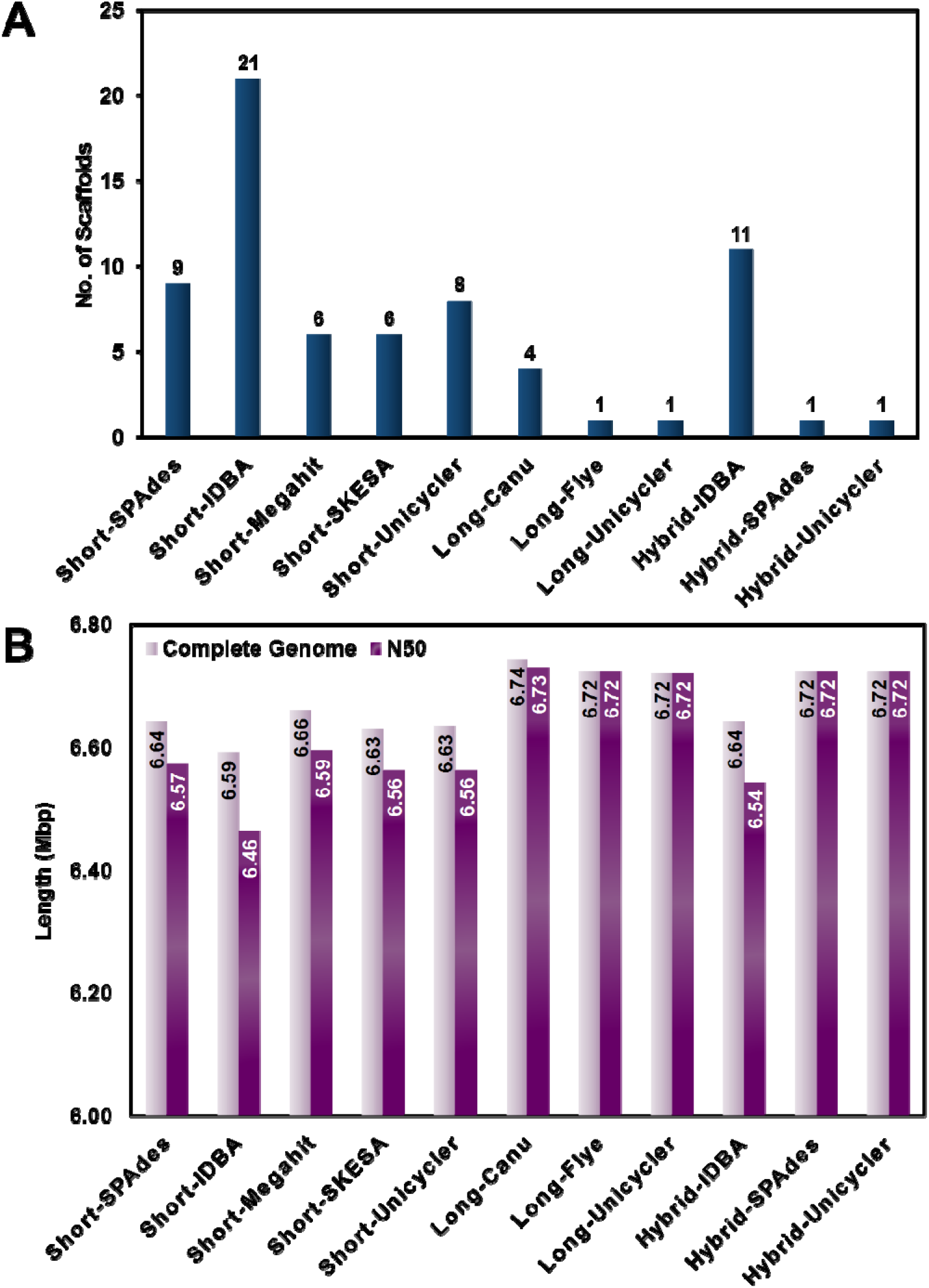
Variations in PPA14 genome contiguity when assembled using different assembly tools. (a) The number of scaffolds generated during the PPA14 genome assembly by different tools (b) The genome size and N50 value of the PPA14 genome assemblies produced by each tool (Molina-Mora et al. 2020). N50 - length of the shortest contig that covers 50% of the genome size.

**Fig. 6.**
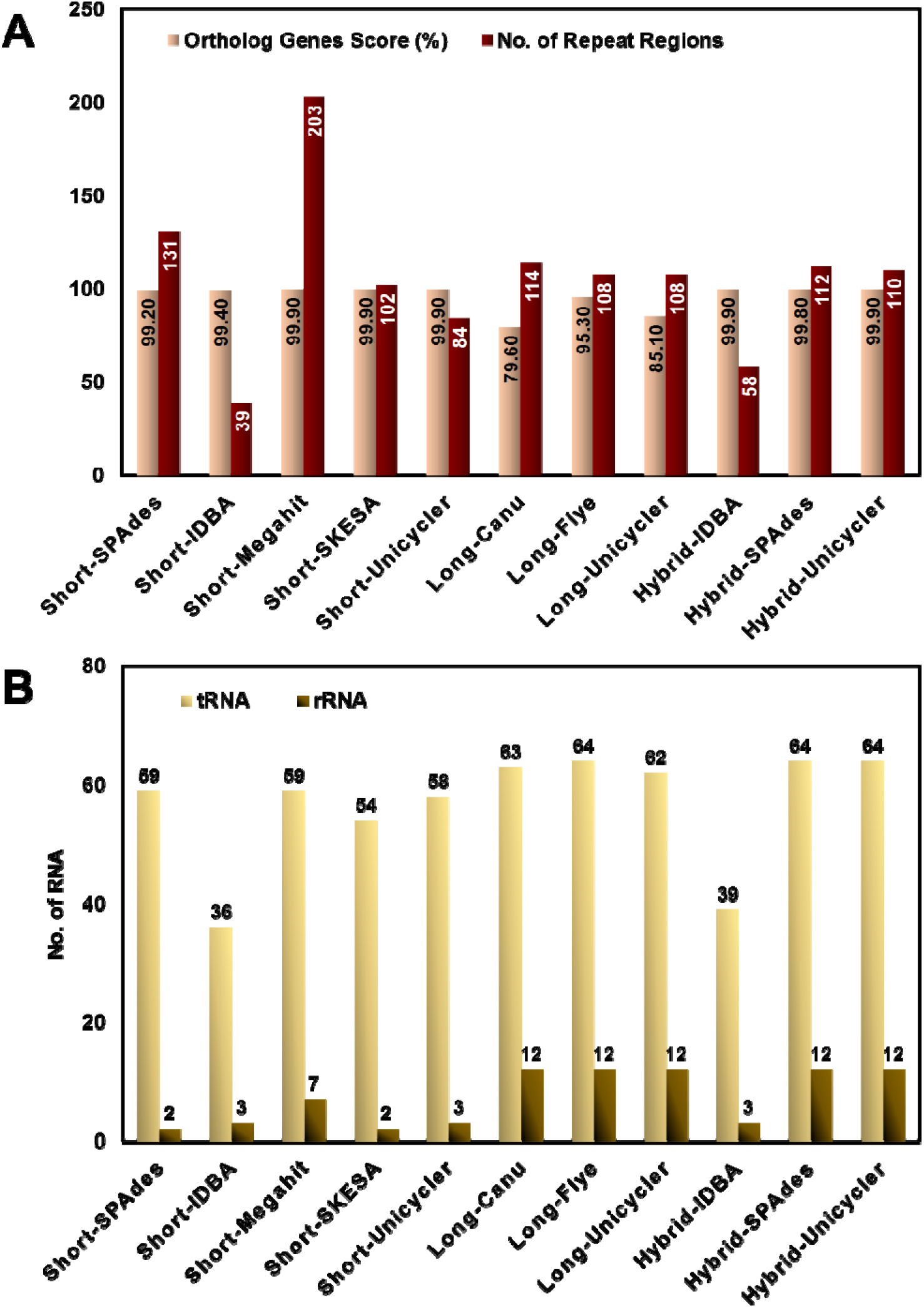
Variations in PPA14 genome completeness when assembled using different tools. (a) Orthologous genes score and the number of repeat regions in different genome assemblies. (b) The number of tRNAs and rRNAs identified in different genome assemblies (Molina-Mora et al. 2020).

Overall, the Hybrid-Unicycler assembly had the best scores based on all the tested parameters and was selected for downstream analysis.

### Prediction of gene functions in the PPA14 genome

The size of the PPA14 genome was 6.72 million base pairs (Mbp) carrying 6315 ORFs. The predicted ORFs were individually analyzed to determine their functional motifs. In addition to the Prokka and RAST-based gene annotations, each ORF was individually evaluated based on previous records to avoid misrepresenting the gene products. Manual curation aims to offer high-quality genome annotations by correcting the inaccuracies in the software outputs. Missing genes, start codon misalignments, and disrupted genes were manually evaluated and fixed. Gene functions were predicted based on the homologous sequences in *P. aeruginosa* genomes that were previously characterized and deposited in reference databases. Out of 6315 open reading frames in the PPA14 genome, functions were predicted for 5042 genes. The remaining 20% of the genome was scored hypothetical as those genes did not have identifiable functions. The PPA14 genome had 64 tRNA, and 12 rRNA genes that comprised of four sets of 23S, 5S, and 16S subunits. It was also observed that the origin of replication or chromosomal replication initiator protein DnaA was located at 483 bp in the PPA14 genome. This gene was followed by DNA polymerase III beta subunit, DNA recombination, and repair protein RecF, DNA gyrase subunit B, and Putative helicase. The order of these genes was similar to the reference *P. aeruginosa* genomes such as PAO1, PA14, PAK, and PA7 (https://pseudomonas.com/). This further validates the gene annotation process carried out in the current study.

### Genomic islands harbored by the PPA14 genome

GI refers to the cluster of genes that have been potentially acquired by a bacterium through HGT. Such gene clusters play a significant role in bacterial genome evolution. In the current study, IslandPath-DIMOB, SIGI-HMM, and IslandPick tools collectively identified 83 GIs in the PPA14 genome. The largest GI detected was 48 kilobase pairs (Kbp) long constituting 54 ORFs. It was observed that 34 out of 83 GIs were longer than 10 Kbp. In total, over 982 genes of horizontal origins were detected in these islands. Linear maps representing the functional motifs of the genes harbored in each GI were created for all the 83 predicted islands. Our results show that nearly 15% of the PPA14 genome carries genes that were acquired through HGT.

### Genetic determinants of the PPA14 antibiotic resistance

A list of 49 ABR genes was identified in the PPA14 genome based on the reference databases, NCBI AMRFinderPlus, MEGARes, CARD, and ResFinder (Table 2). The PPA14 genome carries a set of *aph* genes that code for aminoglycoside O-phosphotransferase conferring resistance against kanamycin, gentamycin, and streptomycin. The genes (*blaPDC*-374 and *blaOXA*-50) that code for class C and D beta-lactamases, respectively conferring resistance against cephalosporins and carbapenems were also identified. The PPA14 genome also had a fosfomycin resistance gene *foxA* coding for FosA family glutathione transferase, and a chloramphenicol resistance gene, *catB7*, coding for chloramphenicol O-acetyltransferase. Additionally, *crpP* coding for a ciprofloxacin resistance protein was also found. Apart from these ABR genes that confer direct resistance to a specific antibiotic, 35 genes that code for multi-drug efflux pumps were detected in the PPA14 genome. This includes the resistance-nodulation-division (RND) superfamily efflux pumps, *mex, mux, tri,* and *opr* that could eliminate a wide spectrum of antibiotics. Moreover, the major facilitator superfamily (MFS), the multidrug and toxic compound extrusion (MATE), and the small multidrug resistance (SMR) family pumps that could resist aminoglycosides, bicyclomycin, and fluoroquinolones were also found. Additionally, the position of 49 ABR genes and 83 genomic islands were located in the PPA14 genome map and visualized using the CGView (Fig. 7). It was found that two sets of *aph(6’)-Id*, *aph(3”)-Ib* genes, *mexXY,* and *crpP* were present within three genomic islands.

**Fig. 7.**
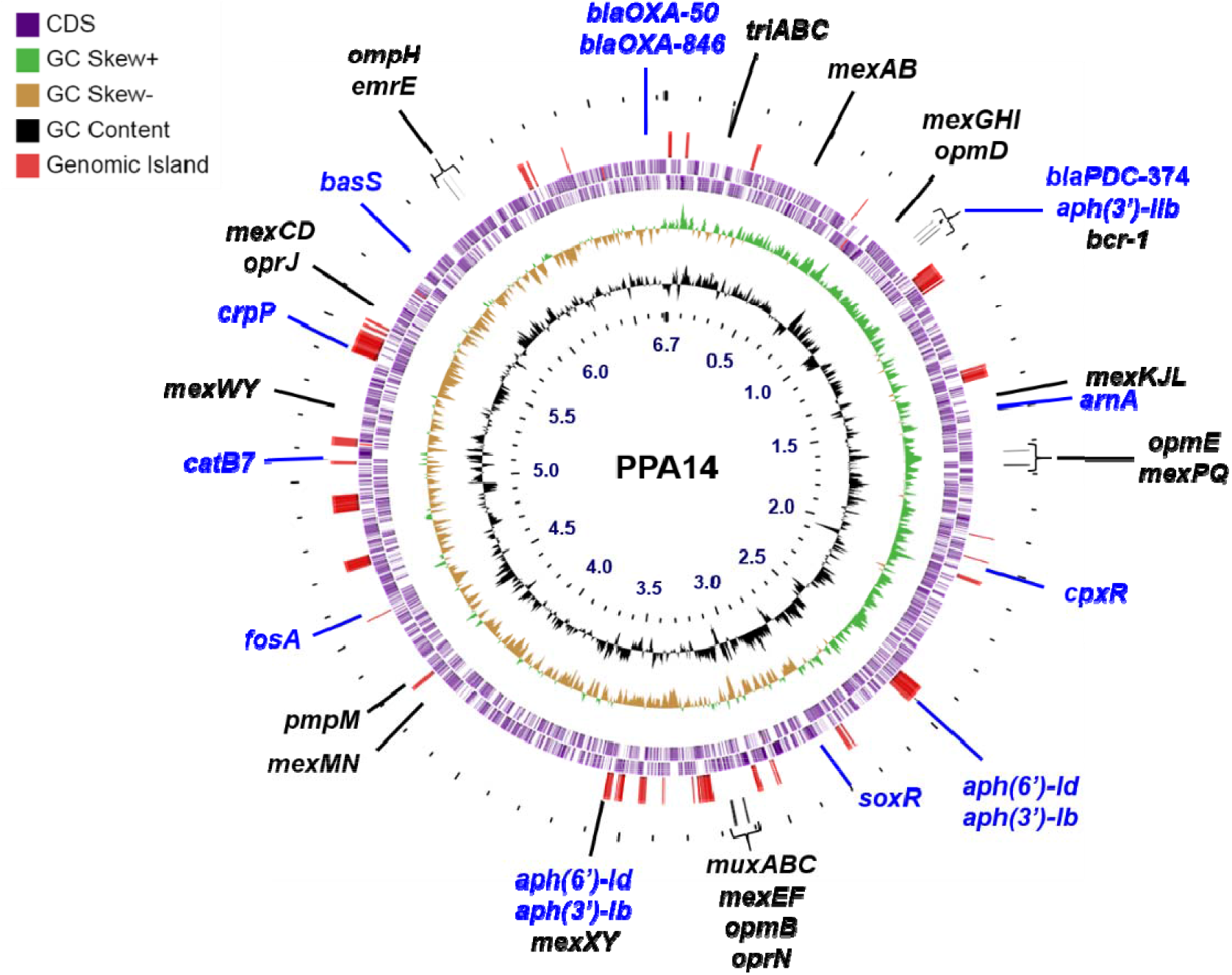
Genome map. The PPA14 genome map representing the AMR genes and genomic islands. The genes in blue font code for proteins that confer direct resistance against specific antibiotics while the genes in black font code for proteins that act as efflux pumps against a wide spectrum of drugs.

**Table 2.**
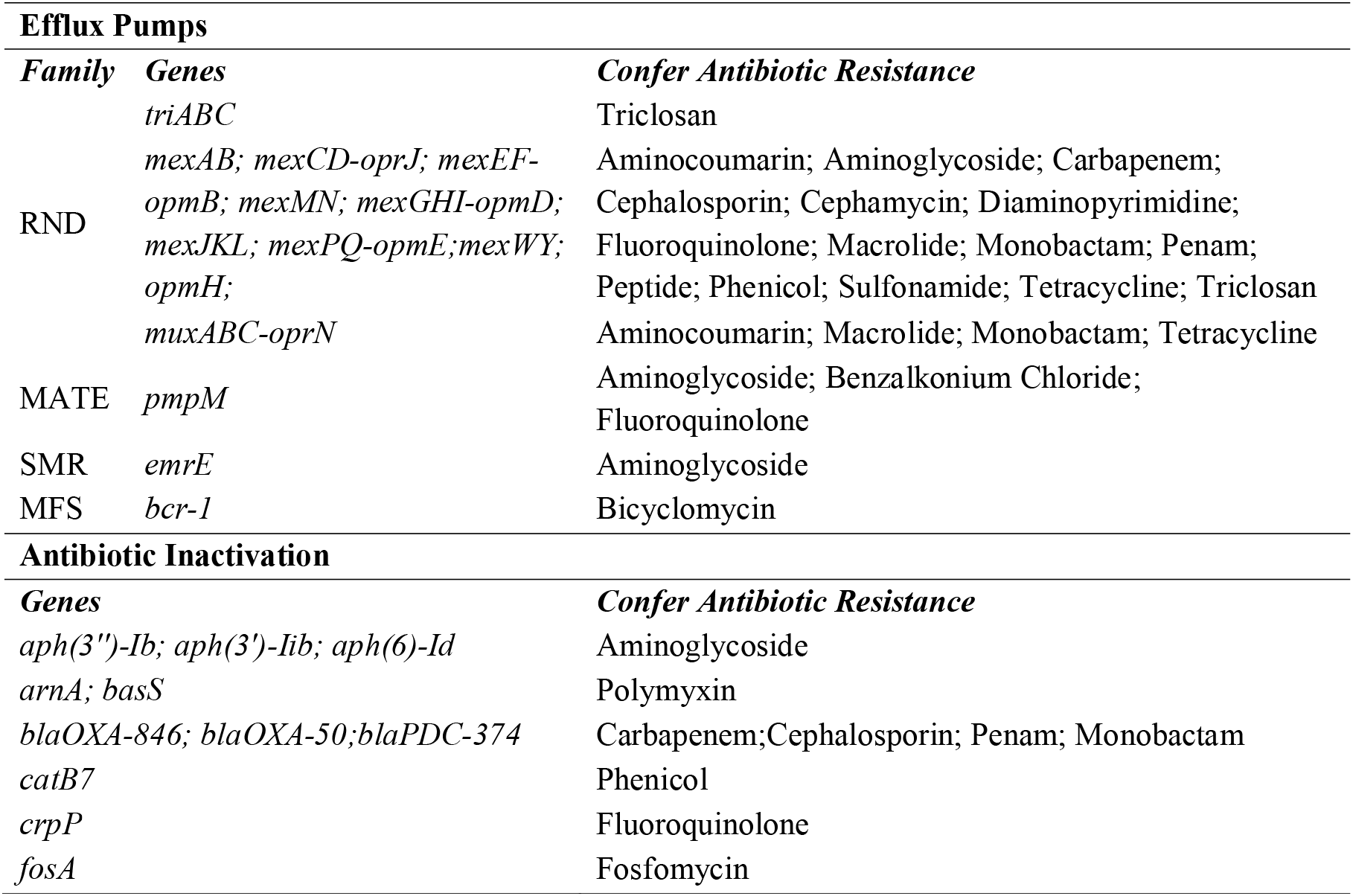
List of antibiotic resistance predicted in the PPA14 genome.

### Genetic determinants of the PPA14 virulence

The virulence-related genes in the PPA14 genome were identified based on the VFDB database (68). More than 200 genes that could confer virulence to the *P. aeruginosa* PPA14 were detected (Table 3). The potential products of these genes included the major virulence factors such as phenazines, rhamnolipids, siderophores (pyochelin, pyoverdine), alginate, lipopolysaccharides, flagellar, pili-related, and secretion system proteins, and exotoxins (Table 3). In addition, the PPA14 genome had a list of genes that could produce lytic enzymes such as protease, lipase, elastase, alkaline and serine proteinase, hemolytic phospholipase, and peptidoglycan hydrolase. Chemotaxis, motility-associated, and ABC transporter genes were also detected in the PPA14 genome (Table 3).

**Table 3.**
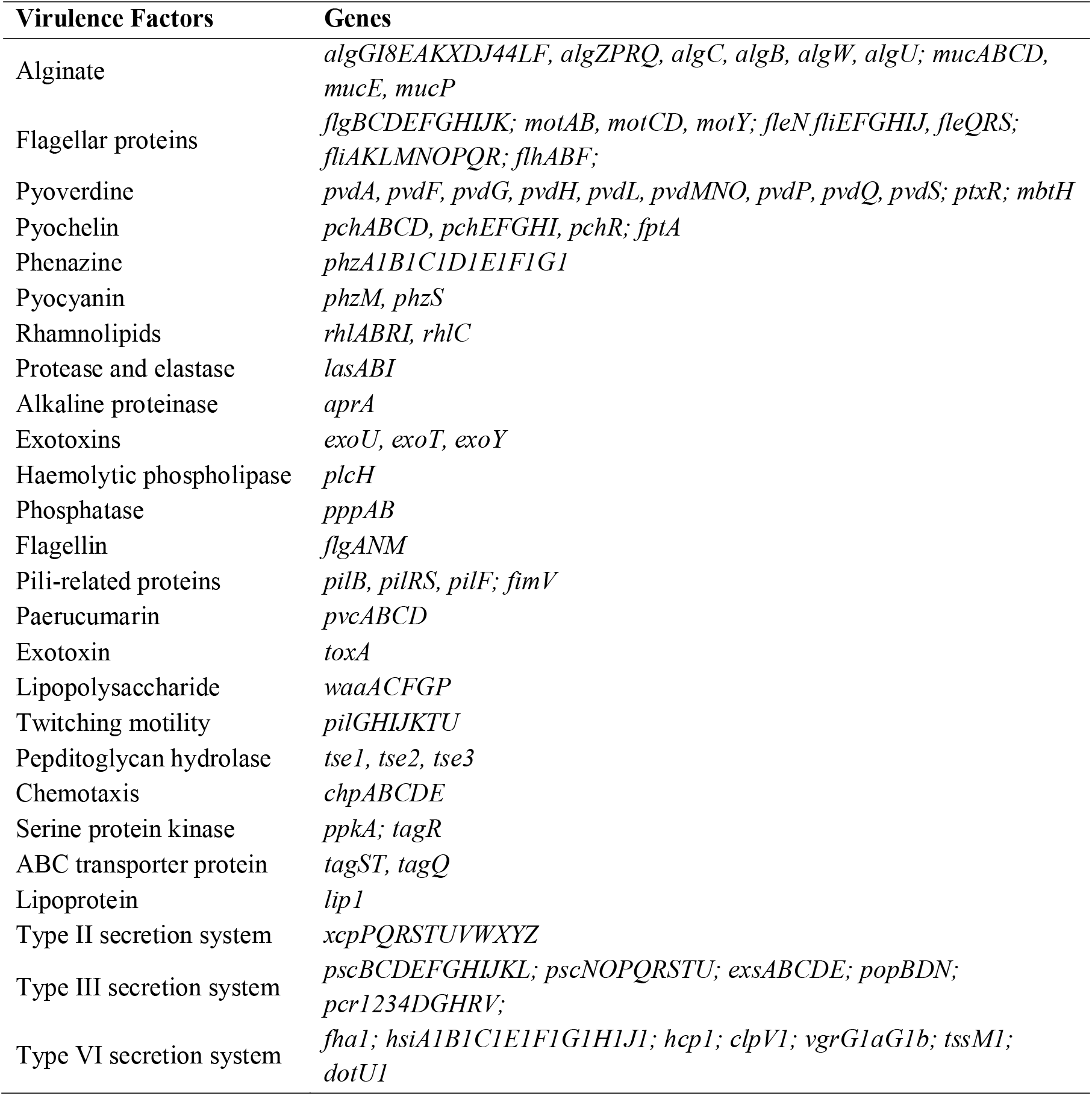
Virulence-related genes in plant-associated *Pseudomonas aeruginosa* PPA14.

### Genomic plasticity in plant-associated and environmental *P. aeruginosa* strains

The complete genomes of *P. aeruginosa* strains that were previously isolated and sequenced from plants and the environment (Table 1) were compared with the PPA14 genome. The *P. aeruginosa* reference strains, PAO1 and PA14, were used as outliers. These reference strains had 6.2-6.5 million base pairs (Mbp) genome size (Fig. 8). The genomes of the plant-associated strains (CR1, YB01, F9676, M18, CFSAN084950, and PPA14) ranged from 6.1 to 6.7 Mbp while the environmental strains (N17-1, RD1-3, YL84, IMP66, IMP67, IMP68, PA1RG, MTB-1, KRP1, B10W, JB2, Ocean-1175, Ocean-1155, HS9, FA-HZ1, and DN1) had the genome sizes of 6.3 to 6.9 Mbp. Overall, the genomes of three environmental strains, ocean-1155, ocean-1175, and DN1 were the largest (6.9 Mbp). Among the plant-associated *P. aeruginosa* strains, PPA14, isolated in the current study had the highest genome size (6.7 Mbp).

**Fig. 8.**
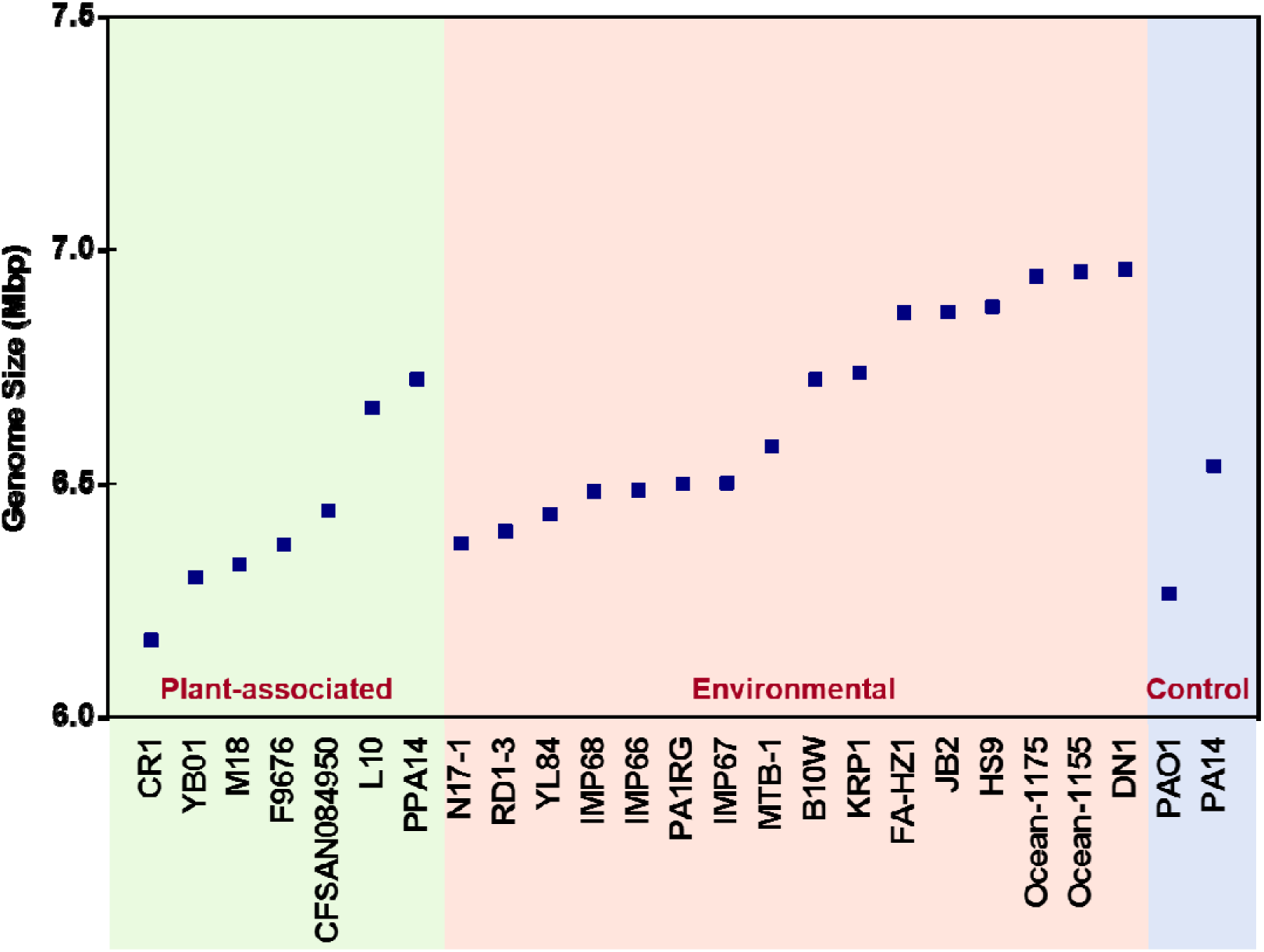
Genomic size of plant-associated, and environmental *Pseudomonas aeruginosa* strains. The graph depicts the strain level variations in the *P. aeruginosa* genome size. The strains are colored based on their source of origin. The plant-associated, environmental, and control strains of *P. aeruginosa* are given green, orange, and blue background colors, respectively.

The total number of protein-coding sequences (CDS), CDS that code for proteins with assigned functions, and non-hypothetical proteins were compared between the plant-associated (CR1, YB01, F9676, M18, CFSAN084950, and PPA14) and environmental strains (N17-1, RD1-3, YL84, IMP66, IMP67, IMP68, PA1RG, MTB-1, KRP1, B10W, JB2, Ocean-1175, Ocean-1155, HS9, FA-HZ1, and DN1) along with the PAO1 and PA14 (Fig. 9). The PAO1 and PA14 genomes had 5.8K and 6.1K CDS, respectively. The number of CDS harbored by the plant-associated *P. aeruginosa* strains varied from 5.7K to 6.2K. The *P. aeruginosa* PPA14 had the highest number of CDS among the plant-associated strains. However, eight out of 16 environmental strains (KRP1, B10W, JB2, Ocean-1175, Ocean-1155, HS9, FA-HZ1, and DN1) had more than 6.2K CDS. The environmental strains with the smallest (N17-1) and largest (DN1) genome sizes had 5.9K and 6.6K CDS, respectively. Nearly 81-83% of the genes in the reference genomes (PAO1, and PA14) coded for proteins with well-known functions. In all of the tested strains, nearly 75-80% of the genes coded for known proteins while the rest were hypothetical.

**Fig. 9.**
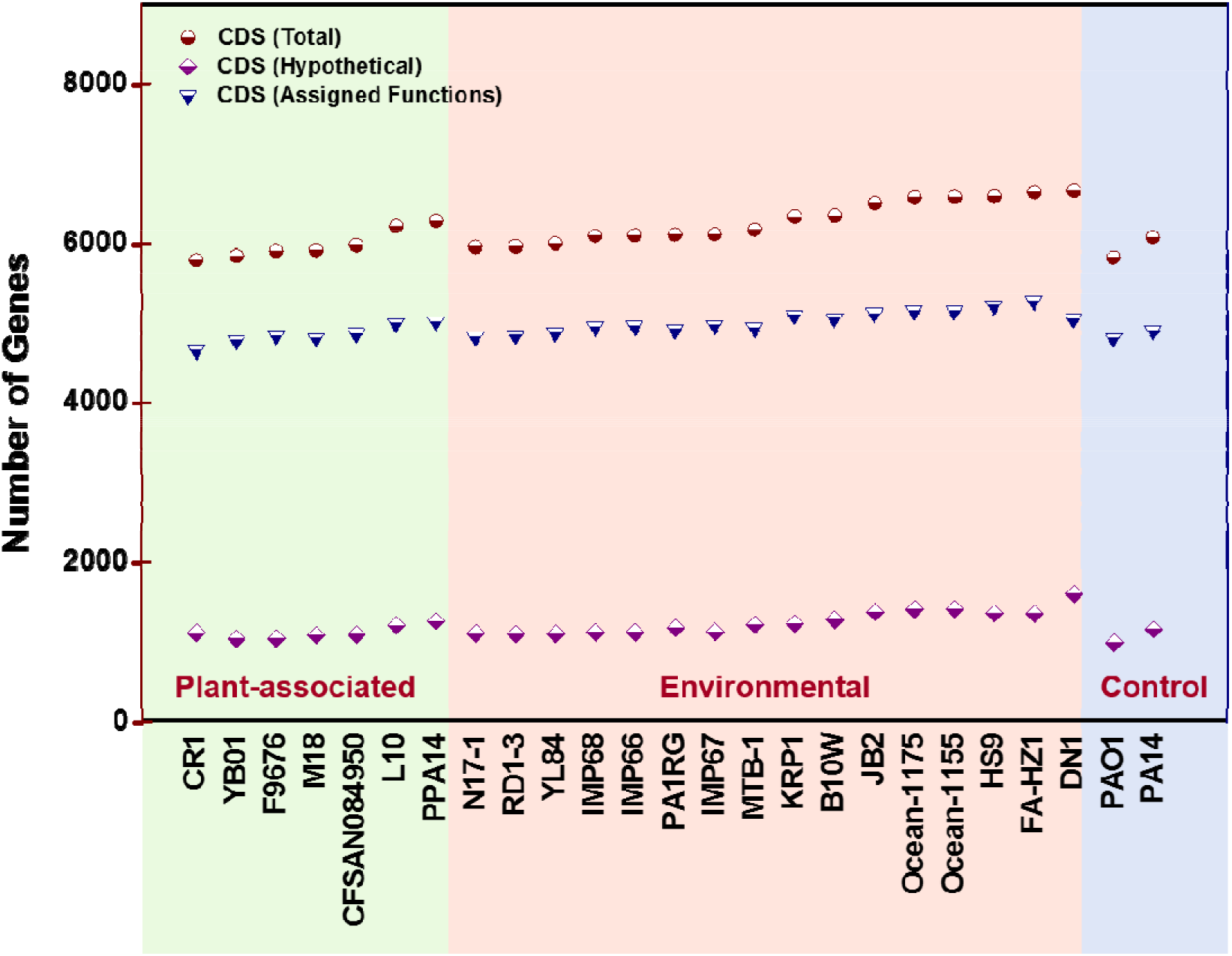
Genomic features of plant-associated, and environmental strains of *Pseudomonas aeruginosa.* The graph represents CDS present in each strain (brown circle), the CDS that codes for proteins with functional assignments (blue triangle), and hypothetical proteins (purple diamond). The plant-associated, environmental, and control strains of *P. aeruginosa* are given green, orange, and blue background colors, respectively.

### Pan-genome profile of plant-associated, and environmental *P. aeruginosa* strains

The number of genes shared across the 23 *P. aeruginosa* strains isolated from the plant (CR1, YB01, F9676, M18, CFSAN084950, and PPA14), and environmental (N17-1, RD1-3, YL84, IMP66, IMP67, IMP68, PA1RG, MTB-1, KRP1, B10W, JB2, Ocean-1175, Ocean-1155, HS9, FA-HZ1, and DN1) ecosystems (Table 1) were estimated using the Roary tool (76). The *P. aeruginosa* reference genomes, PAO1 and PA14, were used as controls (Table 1). The pan-genome is comprised of four components including hardcore, softcore, shell, and cloud genes that were shared by 100%, 95-99%, 15-94%, and less than 15% of the tested genomes, respectively (76). The *P. aeruginosa* genomes compared in this study had 3324 hardcore, 1185 softcore, 1379 shell, and 6736 cloud genes (Fig. 10).

**Fig. 10.**
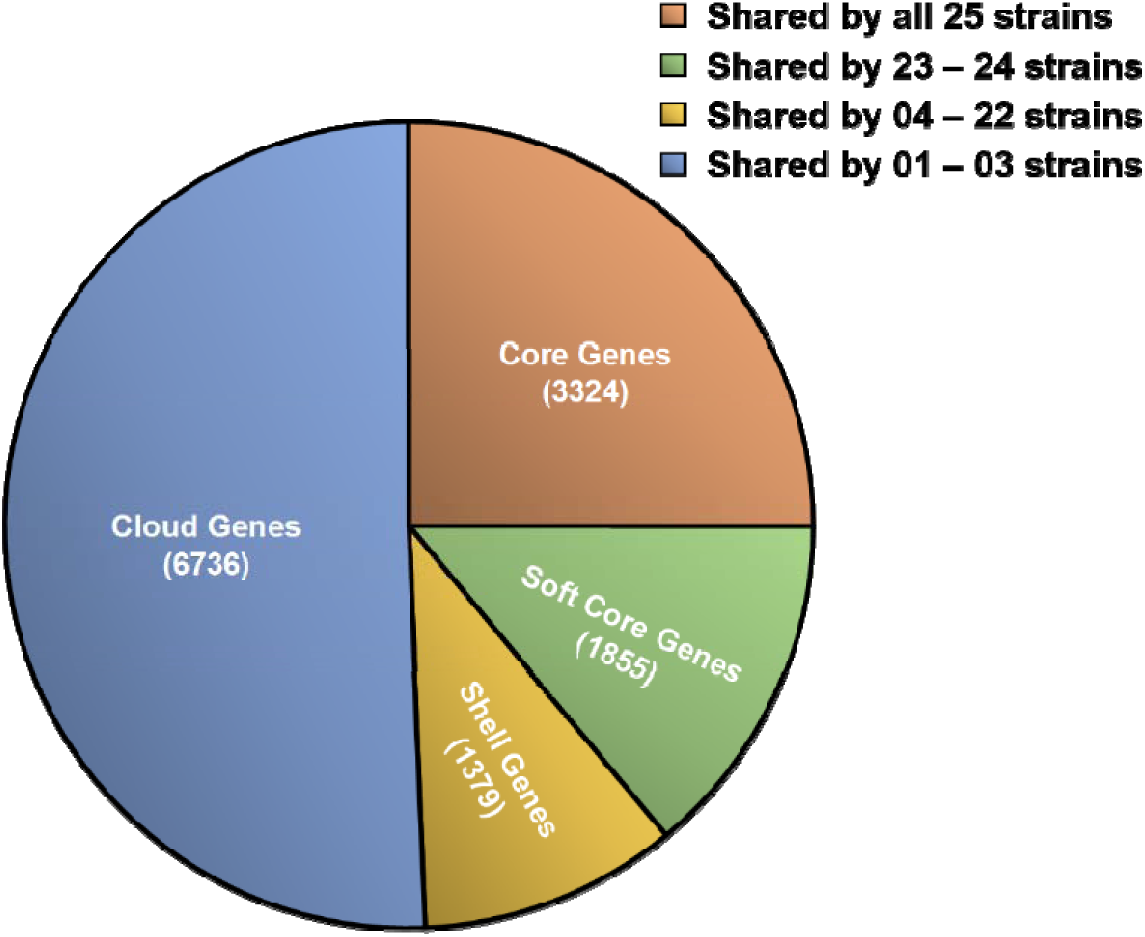
Pan-genome components of *P. aeruginosa* strains. The pie chart represents the number of CDS identified as the hardcore, softcore, shell, and cloud genes while comparing the complete genomes of *P. aeruginosa* strains isolated from plants and environmental ecosystems.

The *P. aeruginosa* pan-genome profiles were visualized using the Phandango interface ((78); Fig. 11). The matrix represented the core and accessory genes identified among the plant-associated, environmental, and reference strains. The hard and softcore were the components of the core genome while the shell and cloud were the components of the accessory genome. Out of 13294 genes identified among the 25 *P. aeruginosa* strains, only 39% occupied the core genome. The remaining 61% were the accessory genes that were absent at least in 22 out of the 25 tested genomes. The number of genes that were unique to each of these *P. aeruginosa* genomes was identified based on the Roary output. The reference strains, PAO1 and PA14, had 53 and 28 unique genes, respectively (Fig. 12). Nearly 65 to 2021 genes were exclusively found in the plant-associated *P. aeruginosa* strains (CR1, YB01, F9676, M18, CFSAN084950, and PPA14). The number of unique genes in the environmental strains (N17-1, RD1-3, YL84, IMP66, IMP67, IMP68, PA1RG, MTB-1, KRP1, B10W, JB2, Ocean-1175, Ocean-1155, HS9, FA-HZ1, and DN1) ranged from 12 to 590. An agricultural *P. aeruginosa*, CR1, isolated from the chili rhizosphere had the highest number of unique genes (2021). An environmental *P. aeruginosa* strain, IMP68 isolated from crude oil had the lowest number (12). The PPA14 genome characterized in the current study had 235 unique genes.

**Fig. 11.**
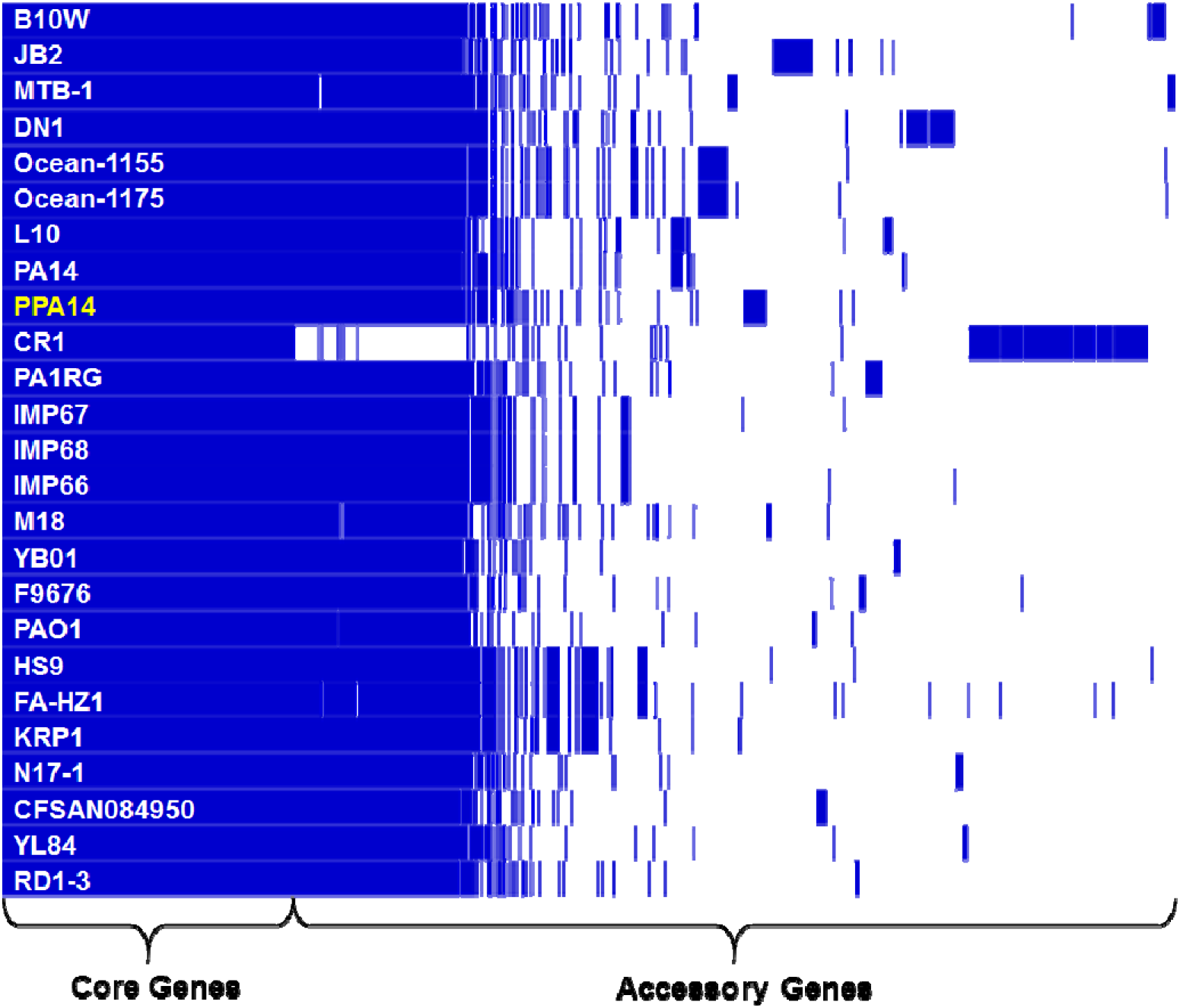
Pan-genome profiles of plant-associated and environmental *Pseudomonas aeruginosa* strains. Visualization of the pan-genome comprising of the core and accessory genes. The *P. aeruginosa*, PPA14, isolated in the current study is highlighted in yellow.

**Fig. 12.**
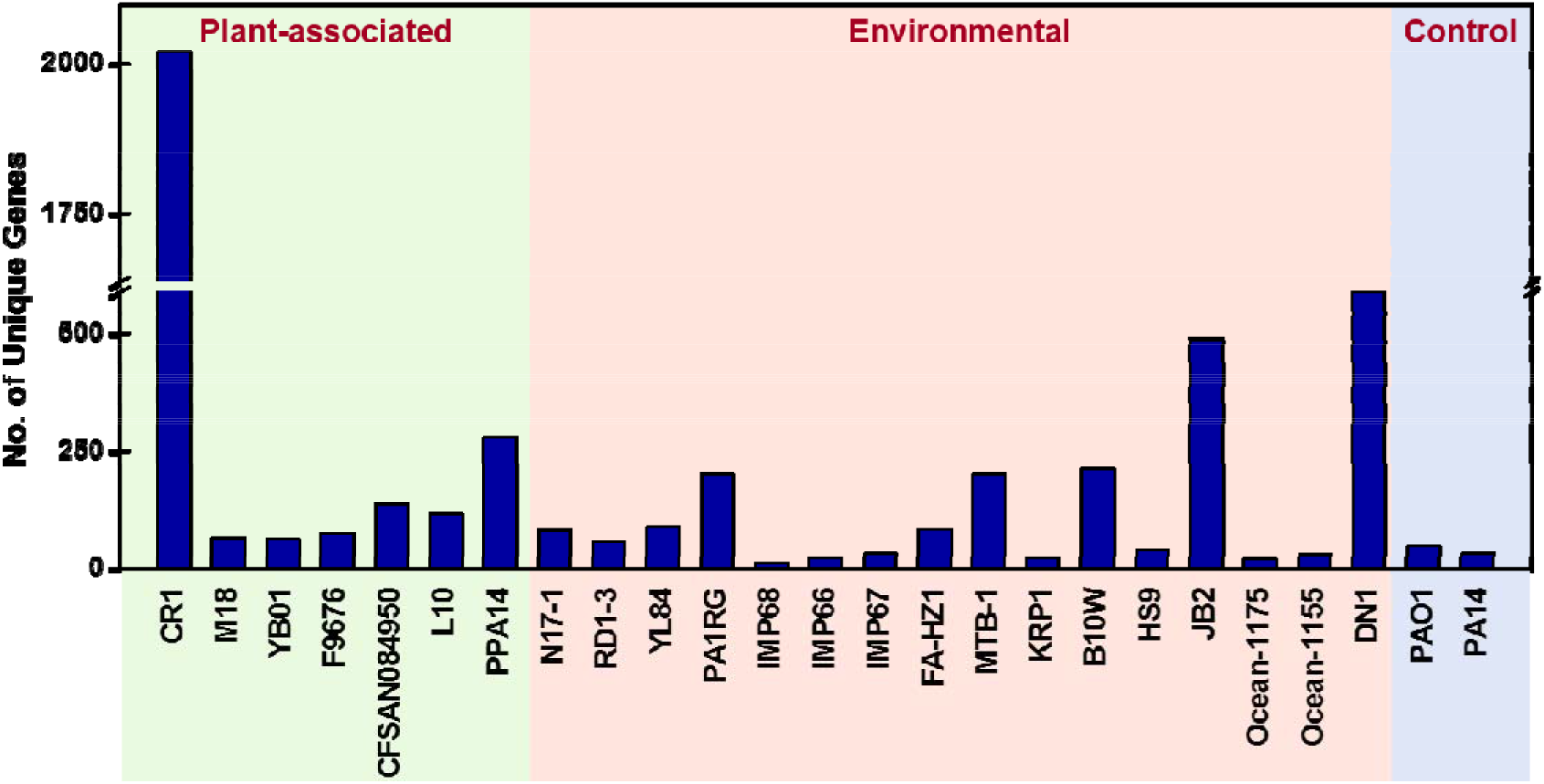
Unique genes harbored by *Pseudomonas aeruginosa* genomes. The graph represents the strain-level variations in the number of unique genes detected in the *P. aeruginosa* genome. The plant-associated, environmental, and control strains of *P. aeruginosa* are given green, orange, and blue background colors, respectively.

**Fig. 13.**
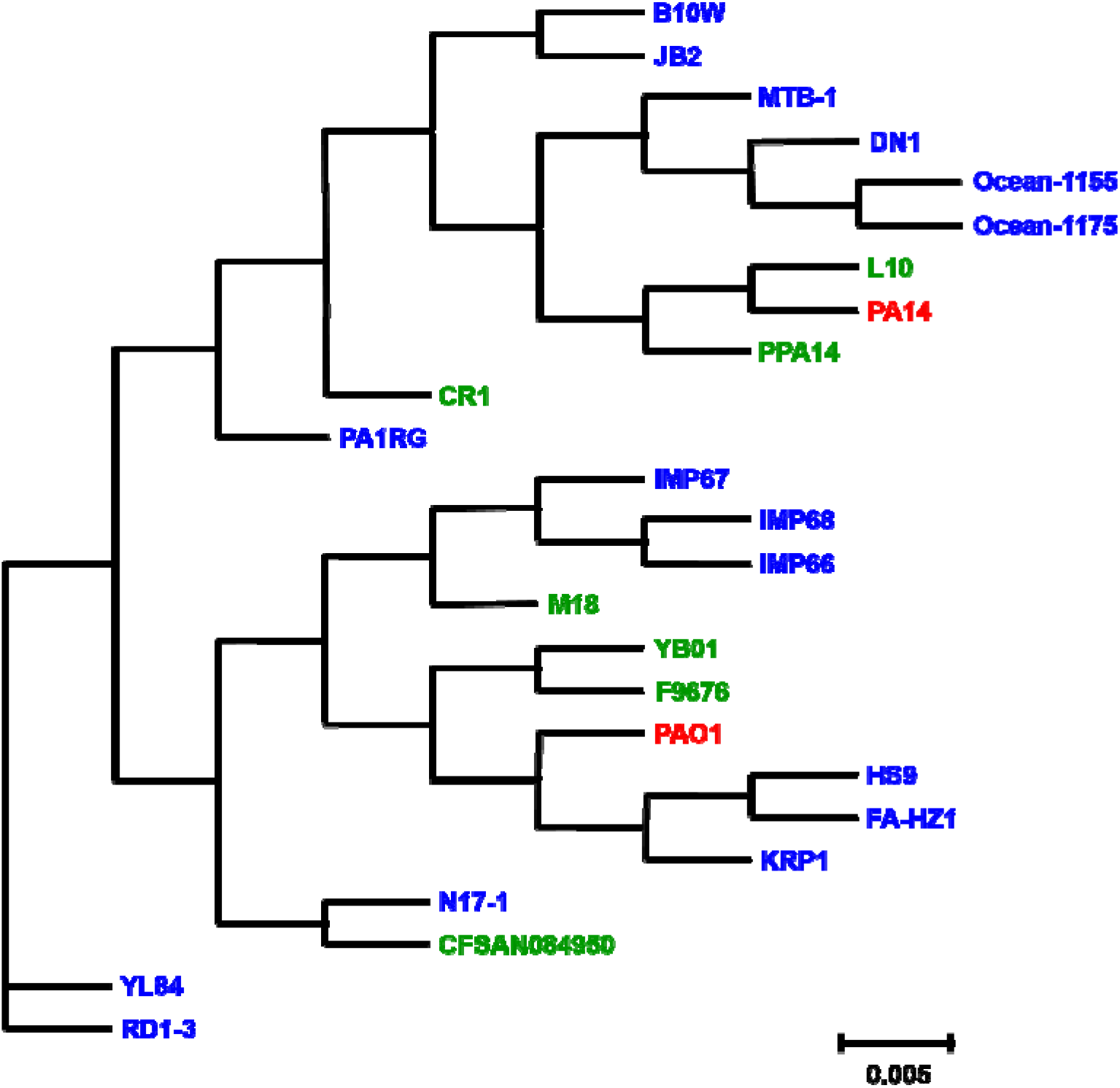
Phylogenetic analyses of *Pseudomonas aeruginosa* strains. The phylogenetic tree was constructed based on the Maximum Composite Likelihood method. The scale value represents the evolutionary distance. The agricultural, environmental, and clinical strains are in green, blue, and red fonts, respectively

### Evolutionary relatedness of plant-associated, environmental, and clinical *P. aeruginosa*

The evolutionary distance between the *P. aeruginosa* strains compared in this study was estimated using the Maximum Composite Likelihood method (80). The phylogenetic analysis clustered the strains based on their pan-genome profile (Fig. 12). The Chili rhizospheric isolate (CR1) and three environmental isolated (PA1RG/hospital sewage; YL84/ compost; RD1-3/ landfill) were the outliers distinct from all the other *P. aeruginosa* strains (Fig. 12). An arugula leaf isolate. CFSAN084950, clustered with N17-1 isolated from farm soil. The rest of the *P. aeruginosa* strains formed two major clades, one with PAO1 and the other with PA14. In the PAO1 clade, three plant-associated (M18/melon rhizosphere; YB01/tobacco stem; F9676/diseased rice sample) and six environmental (IMP66, IMP67, IMP68/crude oil; HS9/soil; FA-HZ1/wastewater; KRP1/methanogenic sludge) *P. aeruginosa* strains were found. The PA14 clade had two plan-associated (PPA14/eggplant rhizosphere; L10/halobiotic reed) and six environmental (B10W/wastewater; JB2/soil; MTB-1/hexachlorocyclohexane contaminated soil; DN1/China soil; Ocean-1155, Ocean-1175/open ocean) *P. aeruginosa* strains.

## Discussion

*P. aeruginosa* is one of the major pathogens involved in hospital-acquired infections and fatalities. Unfortunately, agricultural plants and soil harbor virulent *P. aeruginosa* strains (16, 17) which could be potentially disseminated to humans and animals. In such a case, it is crucial to assess the level of risks associated with the plant-associated *P. aeruginosa*. However, there aren’t many studies on genomic characterization of virulence and pathogenicity in plant-associated *P. aeruginosa*. In the current work, we hypothesized that virulence and drug-resistance genes could be found in the *P. aeruginosa* flourishing in the agricultural ecosystem. To test this hypothesis, we screened the drug-resistance profiles of 18 plant-associated *P. aeruginosa* strains that were previously isolated from cucumber, tomato, eggplant, and chili (16). Among these, an extensively resistant strain PPA14 that shared its virulence factors with the clinical *P. aeruginosa* isolates was selected for genomic analyses. Complete genome analyses of the plant-associated *P. aeruginosa* PPA14 to identify the genetic determinants of its virulence and ABR were the major focus of this paper. Additionally, we have also identified the unique genes in PPA14 and the GIs that were acquired through HGT.

*P. aeruginosa* constantly evolves resistance against multiple antibiotics making itself a Priority Level-I critical pathogen (1). The plant-associated *P. aeruginosa* strains tested in this study exhibited i*n vitro* resistance against cephalosporins, aminoglycosides, macrolides, nitrofurans, tetracyclines, and sulfonamides (Fig. 3a). Nine out of the 18 plant-associated strains (PPA03/cucumber; PPA07, PPA08, PPA10/tomato; PPA13, PPA14/eggplant; PPA16, PPA17, PPA18/chili) had higher resistance than the *P. aeruginosa* PAO1. Drug-resistant *P. aeruginosa* strains have been previously detected in sewage water, effluents, wastewater treatment plants, hydrocarbon-polluted soil, compost, and other contaminated soil and water bodies (8–13, 81). Furthermore, it has been predicted that ABR pathogens get disseminated into the agricultural ecosystem through livestock excreta, manures, and contaminated irrigation systems (82). In specific, the excessive use of antibiotics for the treatment of plant- and animal diseases could lead to the prevalence of ABR pathogens in the agricultural system (83–85). In the present work, we found extensive drug resistance in 50% of the tested plant-associated *P. aeruginosa* (Fig. 3a). Colistin and gentamycin are some of the last resort antibiotics recommended for treating extensively drug-resistant pathogens (86, 87). Even those antibiotics did not control 100% of the agricultural *P. aeruginosa* strains tested in our study (Fig. 3b). Particularly, an eggplant rhizosphere strain, PPA14, was resistant against 85% of the tested antibiotics (Fig. 3a). A detailed phenotypic comparison was done between 18 plant-associated *P. aeruginosa* strains and three clinical isolates (16, 17). Among these, the strains isolated from a wound (PAO1), outer ear infection (ATCC9027), clinical setting (ATCC10145), cucumber endophyte (PPA03), tomato endophytes (PPA08; PPA10), eggplant rhizosphere (PPA13; PPA14), chili rhizosphere (PPA15), and chili endophytes (PPA16; PPA17; PPA18) exhibited shared phenotypic characteristics (Fig. 4). These results highlight the shared virulence and antibiotic resistance between agricultural and clinical *P. aeruginosa* strains. Among the plant-associated *P. aeruginosa* strains that clustered with the clinical isolates, PPA14 was the extensively resistant one and hence was selected for the complete genome analyses.

In the current study, we have detected 49 ABR and 225 virulence-associated genes in the *P. aeruginosa* strain that was isolated from an eggplant rhizosphere (Tables 2 and 3). The environmental occurrence of such ABR pathogens is a globally emerging health concern. Some of the previous studies argue that environmental *P. aeruginosa* strains are highly sensitive to most antibiotics unlike the clinical isolates (88, 89). However, virulent and ABR*P. aeruginosa* strains have been reported in several natural environments like oil-contaminated soil, freshwater springs, domestic sewage, hospital wastewaters, and wastewater treatment lagoons (7, 11, 12, 90–92). Intensive use of antibiotics in poultry farming subsequently leads to the dissemination of ABR *P. aeruginosa* in nearby soil (93). Nearly 75% of the *P. aeruginosa* strains previously isolated from different freshwater resources carried virulence factor genes and exhibited resistance against carbapenems, β-lactams, piperacillin, ceftazidime, and ciprofloxacin antibiotics (94). Also, there have been many reports on plant surfaces and vegetables carrying numerous strains of *P. aeruginosa* (16, 95). More than 200 *P. aeruginosa* strains have been isolated from vegetables collected from different farms and supermarkets (96). However, there were very limited attempts to look into the virulence and ABR determinants in *P. aeruginosa* strains flourishing in farm vegetables. The eggplant-associated strain, PPA14 characterized in the current study harbored genes that could confer resistance against 11 antibiotic classes including penicillins, tetracyclines, cephalosporins, fluoroquinolone, macrolides, sulfonamides, glycopeptides, aminoglycosides, carbapenems, fosfomycin, and monobactam (Table 2; Fig. 7). This strain also possessed 83 GIs that constitutes 15% of its total genetic material. In previously reported *P. aeruginosa* genomes, the genes acquired through HGT have maximumly constituted only to 10% of its genetic material (97–99). This indicates the presence highly versatile and hypermutable genome in the *P. aeruginosa* PPA14 that has acquired a high number of new genetic elements. Moreover, an arsenal of virulence determinants including the phenazines, rhamnolipids, siderophores (pyochelin, pyoverdine), alginate, lipopolysaccharides, flagellar, pili-related, and secretion system proteins, exotoxins, protease, lipase, elastase, alkaline, and serine proteinase, hemolytic phospholipase, peptidoglycan hydrolase, chemotaxis, motility-associated, and ABC transporter genes were also found in the PPA14 genome (Table 3). The incidence of multiple virulence, and ABR genes in the microbes flourishing in agricultural systems could confer alarming threats to human health (100–103).

## Conclusion

The presence of multiple virulence and ABR genes in an eggplant-associated strain reveals the health threats associated with non-clinical *P. aeruginosa*. Furthermore, 12 out of 18 plant-associated *P. aeruginosa* strains characterized in this work exhibited higher *in vitro* resistance than the clinical isolate PAO1. These reports indicate the rising risk of ABR pathogens in agricultural produces. Consumption of raw vegetables (salads) could potentially disseminate ABR pathogens into humans and animals. As it is highly challenging to treat the infections caused by antibiotic-insensitive pathogens the current scenario might escalate the ABR-associated mortality rates in the future. The WHO has already predicted 10 million annual deaths by 2050 due to ABR infections (1). The incidence of ABR pathogens in agricultural produces could be reduced in the future by preventing the (i) excessive use of antibiotics on farm animals (ii) dumping of pesticides on farmlands and (iii) use of sewage-contaminated water for irrigation.

## Abbreviations

ABR: Antibiotic resistance
MDR: Multi-drug resistant
HGT: Horizontal gene transfer
PPA: plant-associated *Pseudomonas aeruginosa*
LB: Luria Bertani
CTAB: hexadecyl-trimethyl ammonium bromide
Prokka: rapid prokaryotic genome annotation
RAST: Rapid annotation using subsystem technology
NCBI: National Center for Biotechnology Information
CARD: Comprehensive antibiotic resistance database
CGE: Center for Genomic Epidemiology
VFDB: Virulence factor database
GI: genomic island
CGView: Circular genome viewer
ORF: Open reading frames
RAxML: Randomized axelerated maximum likelihood
ANOVA: Analysis of variance
Mbp: Million base pairs
RND: Resistance-nodulation-division
MFS: Major facilitator superfamily
MATE: Multidrug and toxic compound extrusion
SMR: Small multidrug resistance
CDS: protein-coding sequences

## Data Summary

All sequence data generated in this study were deposited in NCBI GenBank (Accession no. MT734694 to MT734711).

## Author Contributions

The genomic experiments were conceived and designed by SA and KM. Phenotypic experiments were designed by SA and DB. All the experiments in this paper were performed by SA unless specified otherwise. Whole genome assembly, manual curation, and identification of genomic islands were done by TC, CC, and GP, respectively. Critical analyses of the data and manuscript preparation were done by SA and KM. Finally, all the authors were involved in the critical review of the paper.

## Acknowledgment

Fulbright Doctoral Nehru Research Fellowship awarded to SA by the US Department of State’s Bureau of Educational and Cultural Affairs, and the United-States India Educational Foundation is gratefully acknowledged.

## Ethical statement

The work did not involve humans or experimental animals.

